# Can Folding Models Tell Binders from Bluffers? Evidence from POISK: The Patent-Derived Antibody Dataset

**DOI:** 10.64898/2026.10.24.761513

**Authors:** Daria Tupikina, Andrea Roncoli, Alexander Bujotzek, Brennan Abanades

**Affiliations:** Large Molecule Research, Roche Pharma Research and Early Development, Roche Innovation Center Munich, Penzberg, Germany; Prescient Design – AI for Drug Discovery, Computational Science Center Of Excellence, Roche, Basel, Switzerland; Utrecht University, Faculty of Science – Chemistry, Bijvoet Centre, the Netherlands

**Author notes:** 40th Conference on Neural Information Processing Systems (NeurIPS 2026). Track on Evaluations and Datasets.

## Abstract

Structure prediction models are increasingly deployed as zero-shot digital screens in antibody design, operating under the assumption that high folding confidence implies true binding. However, rigorous validation of this premise has been limited by small test sets, narrow target diversity, and the risk of data leakage from training-adjacent structures. Here we introduce POISK: Patent-extracted Organization of Immunoglobulin Sequence Knowledge, a large-scale dataset derived from the US patent literature, comprising 89,074 antibody-antigen records extracted from 10,110 patents using an LLM-based pipeline. From this resource, we curate a balanced benchmark set of 22,494 complexes: 11,247 patent-validated binders without a known resolved structure paired with putative non-binders constructed from sequence-similar human repertoire antibodies. Using this dataset, we benchmark six leading structure prediction models (AlphaFold-Multimer, Boltz-2, IntelliFold v2, OpenFold3, Protenix-v1, and Protenix-v2) on their ability to discriminate binders from decoys. The best-performing model, Protenix-v2, achieves 72% pairwise accuracy over all pairs and 87% on the most confident half of predictions, and correctly identifies the true binder from a pool of 50 candidates in 29% of trials; this represents a substantial enrichment over random selection, yet remains far from reliable single-candidate identification. Structural consensus across models also discriminates binders from decoys, with pairwise accuracy reaching 73.1% when three or more models converge on a pose (*DockQ ≥* 0.23, covering around 54.2% of pairs). Epitope localization remains unreliable: the best model contacts annotated epitope residues in only 62.8% of cases, indicating that binding discrimination does not establish accurate epitope localization. Our results suggest that current folding models provide useful but limited signal for antibody screening and should be complemented by orthogonal methods. We release POISK as a community resource for future benchmarking and model training.

## 1 Introduction

The rapid progress of protein structure prediction has transformed how protein-protein interactions are studied [Jumper et al., 2021, Abramson et al., 2024]. Co-folding models such as AlphaFold-Multimer [Evans et al., 2021] have been applied to map large-scale interaction networks [Bryant et al., 2022, Humphreys et al., 2021, Didi et al., 2026] and are now routinely used as zero-shot digital screens in antibody discovery and *de novo* protein design. The underlying assumption is that high structural confidence, as measured by metrics like interface predicted template modelling (ipTM), and folding self-consistency imply true binding [Watson et al., 2023, Pacesa et al., 2025, Ren et al., 2025, Stark et al., 2025].

Despite the widespread adoption of this “confidence-equals-binding” premise, rigorous validation remains scarce. Existing benchmarks rely on the Protein Data Bank (PDB) [Berman et al., 2000] and the Structural Antibody Database (SAbDab) [Dunbar et al., 2014]. Because these are the exact repositories used to train the models, temporal holdout sets have become the standard evaluation strategy [Wohlwend et al., 2025, Smorodina et al., 2026]. Such test sets are typically small, cover limited antigen diversity, and carry a persistent risk of data leakage through homologous sequences in the training data.

To address these limitations, we mine the patent literature, a vast source of experimentally validated antibody-antigen interactions that extends well beyond the target space of public structural databases. Using an LLM-based extraction pipeline, we curate 11,247 novel antibody-antigen pairs from US patents, retaining only antibodies that share less than 95% sequence identity with any antibody in SAbDab. To construct a challenging benchmark, we pair each binder’s antigen with a putative non-binding antibody sourced from the human repertoire (OAS) [Olsen et al., 2022a] or generated via germline reversion. The resulting dataset of 22,494 complexes provides a stringent, SAbDab-filtered test bed for evaluating folding models as binding classifiers.

Using this benchmark, we evaluate six structure prediction models, including AlphaFold-Multimer, Boltz-2 [Passaro et al., 2025], IntelliFold v2 [Qiao et al., 2026], OpenFold3 [Ahdritz et al., 2024], Protenix-v1 [Zhang et al., 2026a], and Protenix-v2 [Zhang et al., 2026b], on their ability to separate binders from sequence-similar decoys. We find that the interface-focused ipSAE (interaction prediction Score from Aligned Errors) metric [Dunbrack Jr, 2025] consistently outperforms ipTM for this task. The best model, Protenix-v2, achieves 72% pairwise accuracy across the full benchmark, rising to 87% when restricted to the 50% of pairs with highest structural confidence, and identifies the true binder from a pool of 50 candidates in 29% of trials. Beyond single-model confidence, cross-model structural consensus provides an independent discriminative signal, reaching 73.1% pairwise accuracy when three or more models predict the same binding pose (*DockQ ≥* 0.23), covering around 54.2% of pairs. However, epitope localization remains unreliable, with the best model contacting annotated epitope residues in only 62.8% of cases. These results indicate that current folding models provide meaningful but insufficient signal for antibody screening and should be complemented by orthogonal experimental or computational methods.

## 2 Methods

The pipeline consists of four main steps: data extraction, data preparation, folding, and analysis, as schematically described in Figure 1 and detailed below.

**Figure 1:**
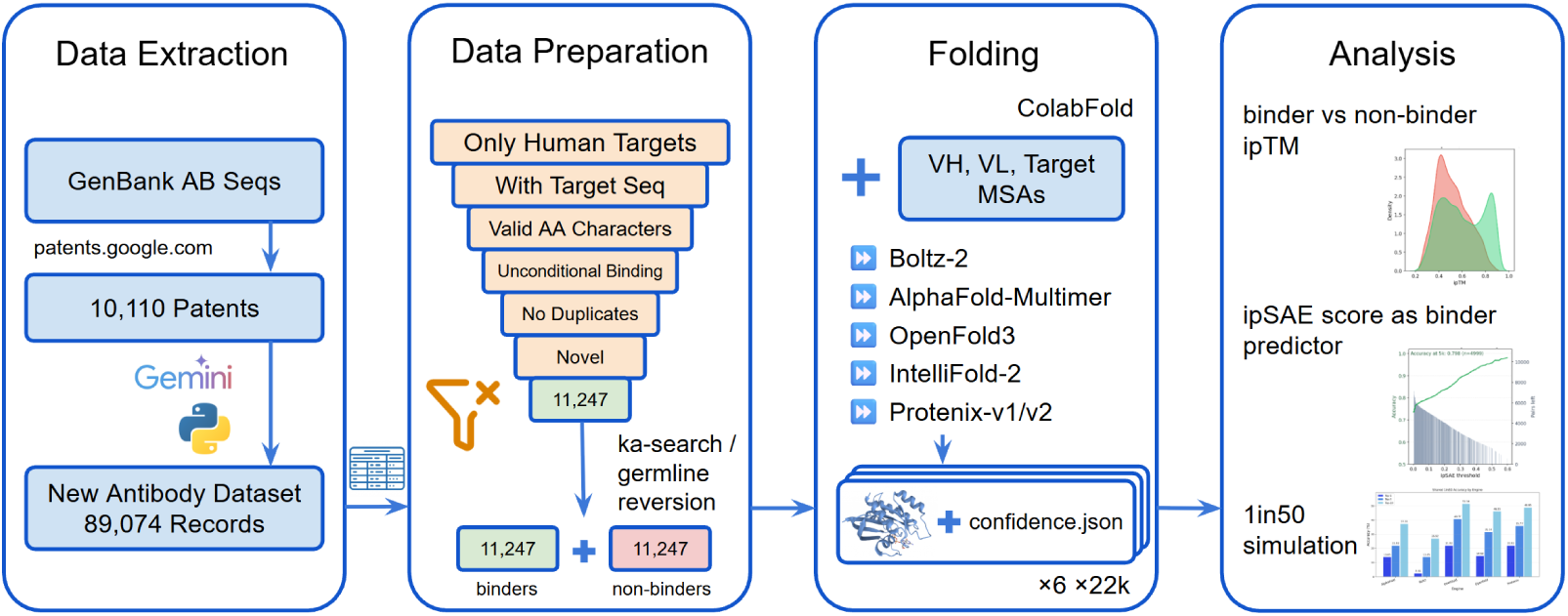
Pipeline overview. Data Extraction: produces an annotated antibody-antigen dataset from the data extracted from GenBank and US patents; Data Preparation: selects complexes eligible for folding and adds putative non-binding antibodies; Folding: precomputes MSAs and runs six folding models; Analysis: computes confidence metrics and benchmarks discrimination power.

### 2.1 Data extraction

We queried GenBank to collect antibody sequences (Heavy, Lambda and Kappa light chains) with lengths between 70 and 1000 amino acids (inspired by [Abanades et al., 2024]). Sequences were classified as antibody chains using ANARCI [Dunbar and Deane, 2016]. In total, we identified 10,110 eligible US patents that contain antibody sequences.

To identify the antigen, pair antibody chains, and extract other relevant antibody details from the unstructured patent text, we developed an extraction pipeline using the Google Generative AI Python library with Gemini 3 Pro model [Anil et al., 2023] and the Pydantic library. We targeted 36 distinct fields, including antibody epitope and experimental binding affinities (see Appendix A for the full list of extracted fields). When the patent did not report an antigen sequence, we searched UniProt using the target identity and organism to retrieve the corresponding sequence [Bateman et al., 2025]. Gemini was also used as an auditor, flagging any information not explicitly mentioned in the patent text. Fields with information flagged by the auditor were excluded, and records flagged to have incorrect chain pairing were removed. In total, the pipeline extracted 89,074 records describing 78,174 unique antibodies, forming the POISK dataset.

Because the resource rests on an LLM extraction step, we complemented the automated auditor with a manual audit of 40 records checked field by field against their source patents: 19 of 20 benchmark records (95%) and 18 of 20 records drawn from the full dataset were correct in every populated field, with 181 of 184 audited field instances correct overall. Two of the three errors were omissions in the conservative direction, a field present in the patent but recorded as absent; the third was a human target labelled *Humanized* rather than *Human*. No record in either sample asserted an antibody–antigen binding pair that its patent does not support. Although no unsupported binding pairs were observed in either sample, this audit does not completely rule out their existence elsewhere in the dataset. More details on this manual audit are given in Appendix B. The extraction and auditor prompts are released alongside the dataset.

### 2.2 Co-folding benchmark data preparation

For benchmarking folding models, we selected a subset of the retrieved data focusing on novel, human-targeting antibodies with explicitly defined antigen sequences. Records indicating conditional (e.g., pH-dependent) binding were excluded because folding models do not yet account for these conditions. Antigen sequence length was restricted to the range of [50, 750] for computational feasibility and duplicate entries were removed. We used Diamond BLASTP [Buchfink et al., 2015] to exclude antibody sequences sharing *≥* 95% sequence identity with any antibody in SAbDab.

After filtering, we were left with a collection of 11,247 unique sets of antibody-antigen sequences. To enrich the dataset with putative non-binding instances, we utilized *KA-Search* [Olsen et al., 2023] against the Observed Antibody Space (OAS) to identify the closest *V_H_* CDR sequence match for each binder, pairing these “near-matches” with the original antigen. For records without a close OAS match (32% of records), we reverted germline-encoded regions to their inferred germline sequences and randomized the CDRH3 loops by sampling from AbLang [Olsen et al., 2022b] predictions while maintaining *≥* 85% total identity. The final dataset we used for benchmarking contained 22,494 total records with an equal split between binding and non-binding instances. We note that the non-binding subset consists of antibodies from naive human repertoires paired with human antigens; while the biological context strongly supports the assumption of non-reactivity, these pairs remain putative negatives and have not been experimentally validated for lack of binding. Appendix D sets out the biological argument for why high-affinity false negatives should be rare, and reports a sensitivity analysis bounding the effect that residual mislabelling could have on our conclusions.

### 2.3 Folding and structural scoring

We evaluated six folding models: Boltz-2, IntelliFold v2, AlphaFold-Multimer, OpenFold3, Protenix-v1, and Protenix-v2. All models used MSAs computed with ColabFold [Mirdita et al., 2022] and no templates. For each complex, the diffusion-based models generated one sample at each of five random seeds, whereas AlphaFold-Multimer used the five parameter models available. The highest confidence prediction was selected for further analysis. In total, folding all 22,494 complexes across the six models required approximately 9000 GPU-hours on NVIDIA B200 GPUs; the folding protocol and compute budget are detailed in Appendix L.

We calculated ipSAE (interaction prediction Score from Aligned Errors) for the top-ranked structure of each record across all six models. Calculations were performed using a modified implementation of the ipSAE tool (https://github.com/DunbrackLab/IPSAE), utilizing both the generated 3D structures and their corresponding PAE matrices. The metric was computed as an asymmetric score treating the paired antibody (*V_H_* and *V_L_* concatenated) as a single chain relative to the target (*V_H_* + *V_L_ →* Target). We employed a standard PAE cutoff of 10 Å and a distance cutoff of 10 Å for all ipSAE calculations.

## 3 Results

### 3.1 Dataset

The POISK dataset (89,074 records from 10,110 patents) is publicly available with full field descriptions at https://zenodo.org/records/21775623. The collection spans over 2000 unique target sequences, with 91.8% of records containing explicit target metadata (Figure 2). Canonical monoclonal *V_H_* /*V_L_* antibodies comprise 85.6% of entries, with the remainder including bispecific (7.4%) and single-domain *V_HH_* (3.9%) architectures. A comparison with the Antibody and Nanobody Design Dataset (ANDD) [Wu et al., 2026] found only 716 shared antibody-antigen entries, underscoring the novelty of the data identified through our patent mining effort.

**Figure 2:**
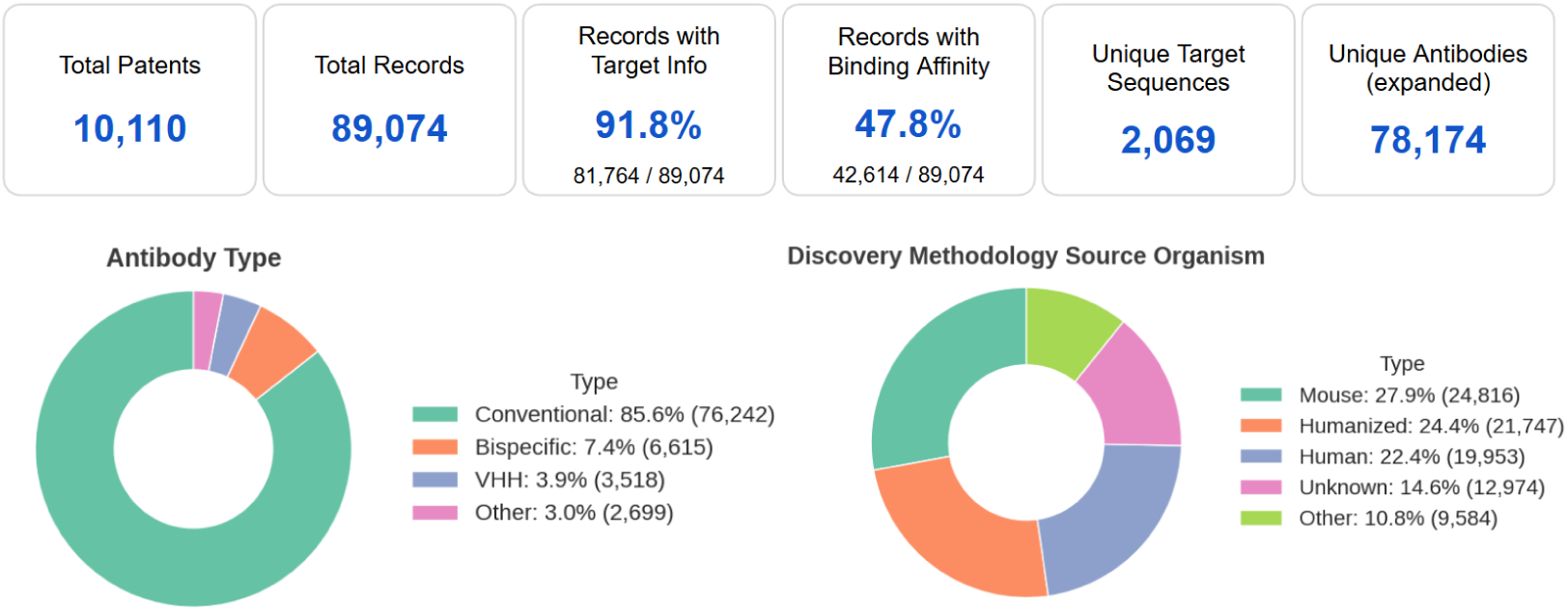
Composition and diversity of the patent-derived dataset. Data overview for 89,074 records extracted from 10,110 patents. The left panel shows antibody architectures, highlighting that conventional *V_H_* and *V_L_* pairings comprise 85.6% of the collection. The right panel displays the discovery methodology source organisms, where mouse-derived and humanized sequences represent the largest subsets.

To evaluate the novelty of our curated folding dataset relative to the structural data typically used to train state-of-the-art folding models, we performed a comparative analysis against the Structural Antibody Database (SAbDab). We first identified structural overlaps by filtering for antigens in SAbDab with *≥* 95% sequence identity to our targets. For this subset, we calculated the minimum Levenshtein edit distance between the concatenated CDR sequences (H1+H2+H3+L1+L2+L3) of our antibodies and those in SAbDab sharing a target. We found that only 1828 of our antibodies targeted antigens with known structures in SAbDab, and only 70 records exhibited a cumulative CDR edit distance of *≤* 10. This minimal structural overlap substantially reduces the risk of data leakage from PDB-derived training data.

### 3.2 Confidence head predicts binding

#### 3.2.1 Baseline performance and metric comparison

We first analysed the distribution of interface predicted template modelling (ipTM) scores to establish a baseline for discrimination power across the six folding engines. Separation between binding and non-binding populations was quantified using Cohen’s *d*, which revealed a clear hierarchy in model performance. Protenix-v2 demonstrated a notable advancement in discriminatory capability, with its Cohen’s *d* increasing from 0.62 in version 1 to 0.75. As illustrated in the density plots (Figure 3), IntelliFold v2 and both Protenix models exhibit clearly distinguishable peaks for the binder populations. This separation suggests a more robust ability to identify true positives compared to the substantial distribution overlaps observed in earlier models.

**Figure 3:**
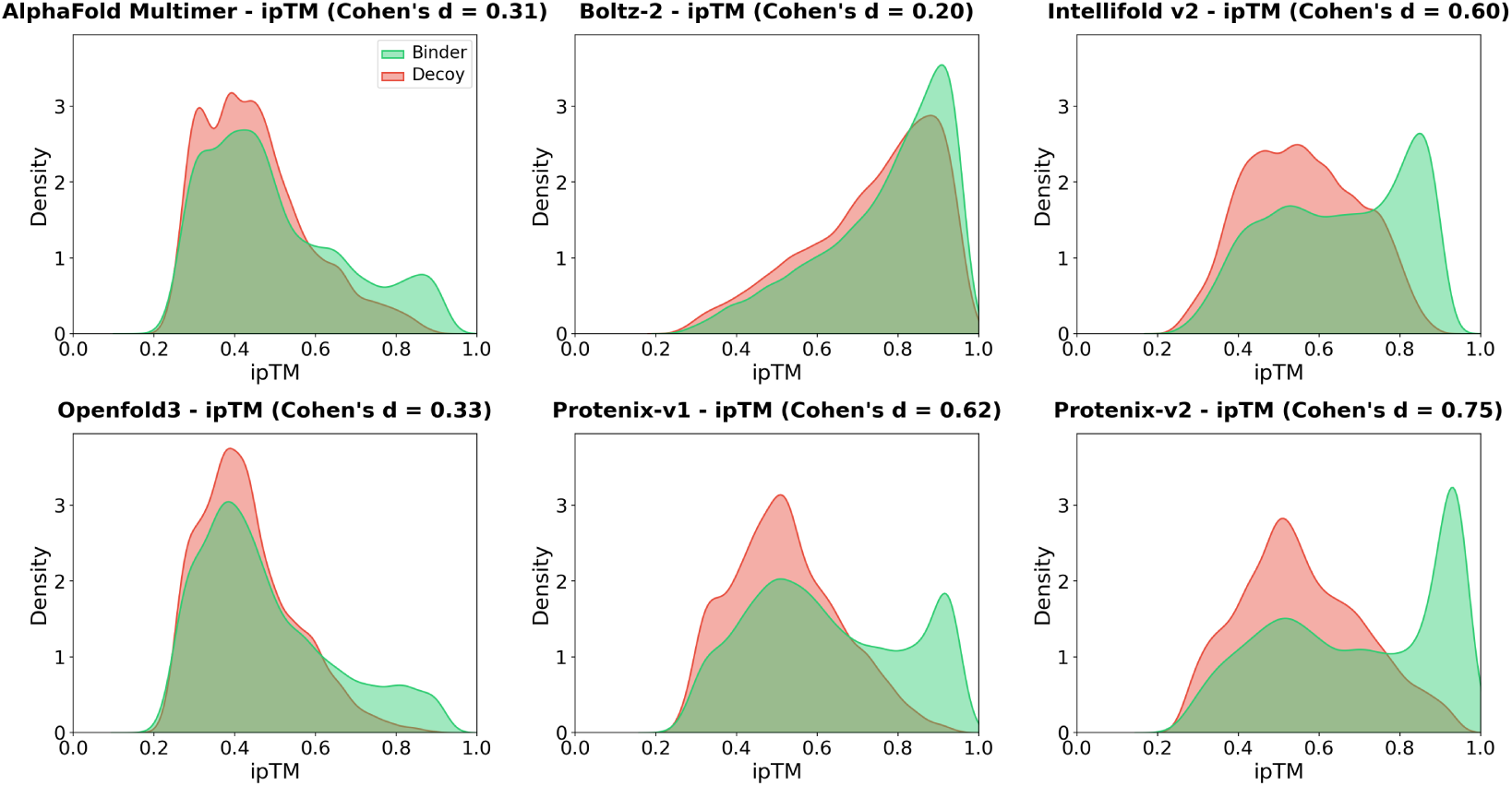
Discrimination performance of six structure prediction models using the ipTM metric. Kernel density estimates illustrate the separation between binder (green) and decoy (red) populations. The discriminatory power of each engine is quantified by Cohen’s *d*, revealing a hierarchy in performance between earlier models and more recent architectures.

We found that alternative metrics, specifically ipSAE described in section 2.3, yielded higher Cohen’s *d* scores than standard ipTM (Table 1). This metric achieves superior class separation, though the values tend to cluster near zero because of the stringent confidence requirements inherent to the calculation. Distribution densities for these scores are provided in Supplementary Figure 8. We selected the full Fv ipSAE as the primary evaluation metric for all subsequent analyses. Unlike ipTM, which scores entire chains and is often confounded by disordered regions or non-interacting accessory domains, ipSAE restricts its calculation to interchain residue pairs below a specific Predicted Aligned Error (PAE) threshold, isolating the interface signal and more effectively separating true binders from background noise [Dunbrack Jr, 2025].

**Table 1:** Summary of benchmark performance for six structure prediction models. Cohen’s *d* values quantify the separation between binder and decoy populations for both ipSAE and ipTM metrics. Pairwise accuracy represents the model’s ability to identify the true binder at the dataset midpoint, where the 50% of pairs with the lowest structural confidence are filtered out. The 1-in-50 Top-1 accuracy indicates the frequency of correctly ranking a single true binder against 49 randomly selected decoys.

| Model | Cohen's *d* |  | Accuracy at | 1-in-50 |
|---|---|---|---|---|
|  | ipTM | ipSAE | 50% Pairs | Top-1 Acc. |
| AlphaFold-Multimer | 0.31 | 0.39 | 0.68 | 0.12 |
| Boltz-2 | 0.20 | 0.23 | 0.60 | 0.03 |
| IntelliFold v2 | 0.60 | 0.66 | 0.80 | 0.25 |
| OpenFold3 | 0.33 | 0.43 | 0.67 | 0.12 |
| Protenix-v1 | 0.62 | 0.71 | 0.80 | 0.23 |
| Protenix-v2 | **0.75** | **0.88** | **0.87** | **0.29** |

#### 3.2.2 ipSAE predictive power on data pairs

Standard *de novo* design relies on the assumption that high-confidence scores effectively distinguish true binders from near-homologue decoys. We tested this assumption by applying ipSAE threshold sweeps to evaluate how structural confidence correlates with predictive accuracy across the six models (Figure 4). These sweeps track the remaining number of binder-decoy pairs as the threshold tightens and the pairs with both ipSAE values below the threshold are filtered out. At each step, we measured how often the model correctly assigned a higher score to the true binder within each pair. We chose the dataset midpoint as a standardized benchmark to compare performance across the different architectures. This point represents the stage where 50% of pairs with the lowest maximum ipSAE are filtered out. Removing these low-confidence pairs separates the model’s ability to rank binders from cases where it failed to produce a confident prediction for either candidate: when both members of a pair score near zero, the comparison reflects folding failures rather than meaningful binding discrimination.

**Figure 4:**
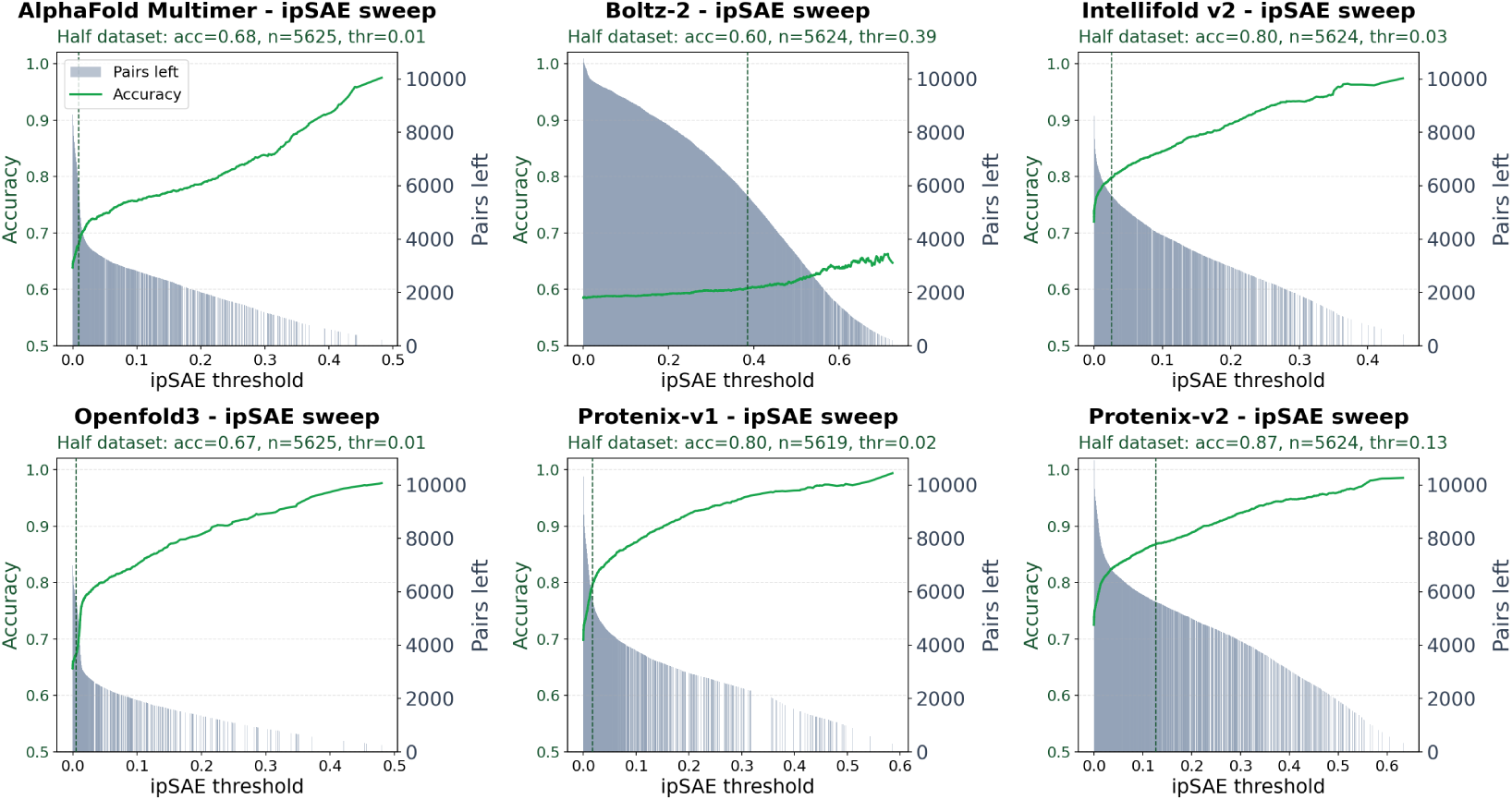
Predictive power of ipSAE across confidence thresholds. The green line tracks the accuracy of identifying the true binder by selecting the higher-scoring candidate in each pair as the ipSAE threshold increases. Gray bars represent the number of pairs remaining in the analysis, as pairs are filtered out if both records score below the threshold.

Under this benchmarking criterion, Protenix-v2 demonstrated the highest selectivity, achieving an accuracy of 0.87 at the half-dataset mark. This result indicates that within the remaining half of the dataset, the candidate with the higher ipSAE correctly identifies the true binder in nearly nine out of ten instances. These findings highlight the utility of the ipSAE as a robust filter for high-confidence structural predictions in antibody discovery. Over all 11,247 pairs, without this filter, Protenix-v2 reaches 0.72, and accuracy rises monotonically as coverage tightens.

The classification performance remains robust across negative data generation strategies: across evaluated engines accuracy for complexes involving OAS KA-Search derived non-binding decoys is comparable to performance on pairs generated through germline reversion (Supplementary Figure 10). While ipSAE scores generally decrease for larger targets across most models, IntelliFold v2 exhibits an unexpected positive correlation between structural confidence and target sequence length (Supplementary Figure 9).

#### 3.2.3 1-in-50 zero-shot screening simulation

While the pairwise sweeps demonstrate the theoretical separation power of the metric, a more realistic discovery scenario involves identifying a single hit within a vast library of non-binding candidates. To evaluate model performance under these conditions, we conducted a “1-in-50” zero-shot digital screening simulation. Unlike the balanced pairwise benchmarks, this test requires the model to correctly rank one true binder against 49 randomly selected non-binding decoys for the same target. We identified 27 targets from the dataset that possessed at least 75 patent-derived binders and performed 20 independent simulation tests per target, resulting in a total of 540 unique discovery trials. Each simulation was repeated over 25 independent random seeds to assess stability. For each trial, we measured the frequency with which the true binder was ranked within the top-1, top-5, and top-10 highest ipSAE scores (Figure 5).

**Figure 5:**
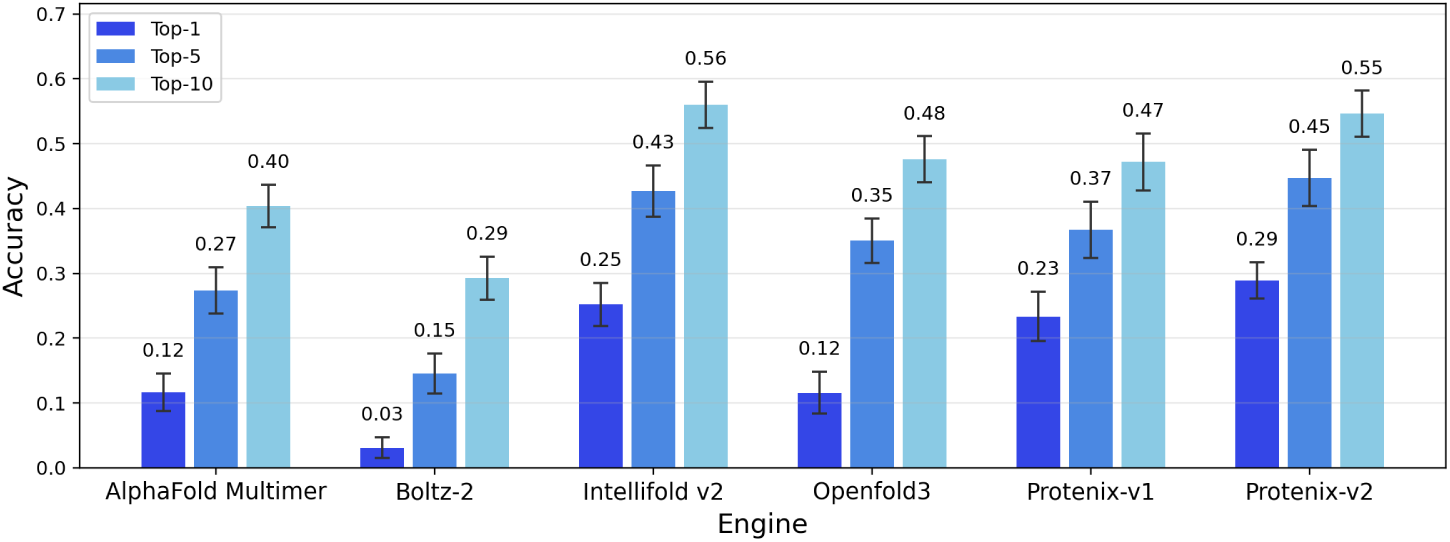
Performance of folding models in a 1-in-50 screening simulation. For each trial, one binder and 49 decoys targeting the same antigen were randomly selected. Bars show mean top-1, top-5, and top-10 accuracy (by ipSAE ranking) over 25 independent runs with different random seeds; error bars indicate *±*2*σ* (sample standard deviation across runs). Results span 540 trials over the 27 targets with at least 75 patent-derived binders.

Protenix-v2 achieved the highest selectivity, identifying the true binder as the top-ranked candidate in 28.9% of tests, a 14-fold enrichment over the 2% random baseline. Protenix-v2 and IntelliFold v2 demonstrated comparable retrieval in broader rankings, placing the true binder in the Top-10 in 54.7% and 56.0% of cases, respectively. In contrast, Boltz-2 reached a Top-1 accuracy of 3.1%, close to the 2% random baseline. These results confirm that ipSAE-based ranking can substantially enrich for true binders in realistic screening scenarios.

#### 3.2.4 Correlation with affinity and structural novelty

We next asked whether structural confidence captures binding strength, a property these models were not explicitly trained on, or is biased by proximity to known structures. We standardized the heterogeneous affinity and potency measurements in our dataset (*K_D_*, *K_A_*, *IC*_50_, *EC*_50_) to pAffinity (*−* log_10_ [M]) across the 5207 binders with reported values. Binned comparisons show a marginal trend toward higher ipSAE at higher affinity, but the signal is too weak to be predictive (Supplementary Figure 11).

For the 1828 records targeting antigens with existing SAbDab entries, we found a similarly weak negative correlation between ipSAE and CDR edit distance (Supplementary Figure 12), suggesting that models assign slightly higher confidence to sequences closer to their training distribution. This correlation is likely weakened by the composition of our dataset: CDR edit distances of *≥* 10 represent functionally distinct antibodies, and the vast majority of our records fall in this regime.

### 3.3 Structural consensus, correctness, and predictive potential

A confidence score conflates two signals: whether a predicted interaction is real and how accurately the model has placed it in 3D space. A low score for a true binder may therefore reflect either a failed fold or a poorly calibrated confidence head, rather than evidence of non-binding. To isolate the binding signal from model-specific scoring artifacts, we next examined the predicted structures directly.

#### 3.3.1 Predicting binding with structural prediction consensus

Just as confidence scores vary across engines, the predicted structures similarly show varying degrees of convergence as illustrated in Figure 6 and Supplementary Figure 15. We hypothesized that geometric agreement across models could serve as a binder predictor independent of the individual confidence heads. To test this, we calculated *DockQ* scores [Basu and Wallner, 2016] for the Fvantigen interface and defined consensus as two or more models predicting the same pose above a given threshold. For any given pair, a prediction was considered successful if the binder demonstrated structural agreement amongst more models than its decoy.

**Figure 6:**
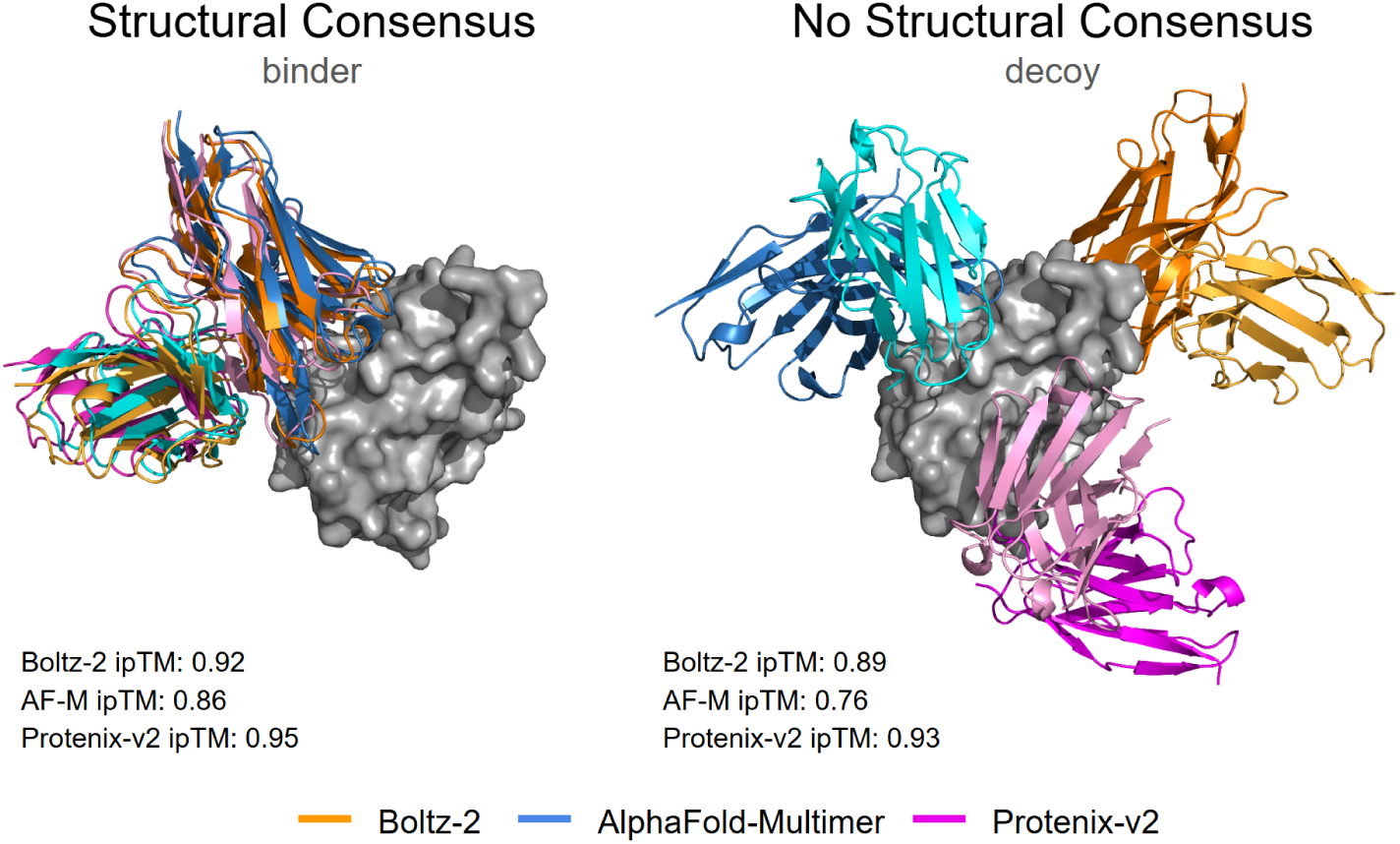
Illustrative examples of cross-model structural agreement and disagreement. Predicted antibody-antigen docks from Boltz-2, AlphaFold-Multimer, and Protenix-v2 are shown for two antibodies targeting the same antigen. Left: when modelling a true binder, the three models converge on a consistent binding pose with high confidence. Right: when modelling a generated decoy, the models predict with high confidence divergent binding poses against the same target for a different antibody.

The results of this consensus approach reveal a strong correlation between geometric convergence and binding. When at least three models agree on a pose at a *DockQ* threshold of 0.23, the pairwise accuracy is 73.1% and covers 54.2% of the pairs. As shown in the sweep plots (Figure 7 and Supplementary Figure 16), requiring more models to agree (*k*) while tightening the *DockQ* threshold substantially improves accuracy. For instance, at *k ≥* 3 and *DockQ ≥* 0.80, accuracy reaches 96.8%. However, this gain in accuracy is offset by a rapid reduction in the number of pairs where models agree, leaving only 3.0% of pairs eligible for classification at this stringent consensus level. While ipSAE remains the more versatile metric for broad library ranking, cross-model consensus provides a complementary signal for prioritizing candidates when multiple folding models strongly agree on a binding pose.

**Figure 7:**
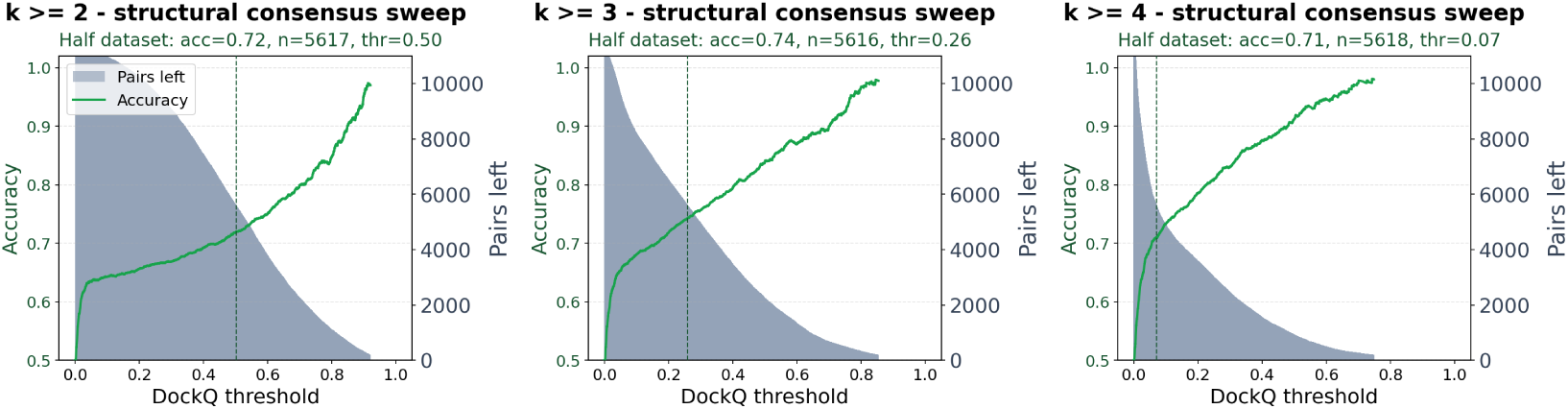
Binding discrimination using cross-model structural consensus for *k ≥* 2, *k ≥* 3, and *k ≥* 4. For each DockQ threshold, bars indicate the number of binder-decoy pairs retained (i.e. where at least *k* models agree on a structure for at least one record in the pair), and the green line tracks pairwise accuracy when classifying the record with more agreeing models as the binder. Increasing both the DockQ threshold and the required number of agreeing models improves accuracy, but at the cost of sharply reduced pair coverage.

#### 3.3.2 Epitope localization and structure prediction correctness

Although we lack experimentally resolved structures to definitively assess prediction correctness, we leveraged the epitope residue annotations from source patents as a proxy to evaluate physical plausibility. These annotations could be mapped to the folded antigen constructs for 1232 binder records in our dataset, allowing us to directly evaluate whether models placed the antibody at the correct binding site. We counted a record as recovered when at least one annotated epitope residue had a heavy atom within 4Å of an antibody heavy atom (Table 2). Even the best-scoring model, Protenix-v2, placed the antibody in contact with the annotated epitope in only 62.8% of cases, indicating that the accurate folding of antibody-antigen complexes remains a substantial challenge for current structure prediction models.

**Table 2:**
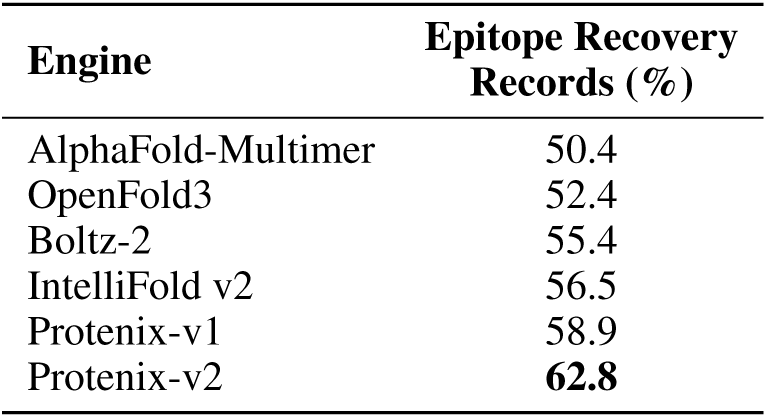
Epitope Recovery Rate by Engine.

| **Engine** | **Epitope Recovery Records (%)** |
|---|---|
| AlphaFold-Multimer | 50.4 |
| OpenFold3 | 52.4 |
| Boltz-2 | 55.4 |
| IntelliFold v2 | 56.5 |
| Protenix-v1 | 58.9 |
| Protenix-v2 | **62.8** |

## 4 Discussion

Structure prediction models can discriminate antibody binders from sequence-similar decoys well above chance (Protenix-v2 achieves 87% pairwise accuracy when restricted to the 50% of pairs with highest structural confidence), but the gap between statistical discrimination and reliable hit identification remains large. In the 1-in-50 screening simulation, even the best model ranks the true binder first in only 29% of trials. This is a substantial enrichment over the 2% random baseline, and likely sufficient to add value as one stage of a multi-step screening funnel, but insufficient to replace experimental validation or to be used as the sole criterion for candidate selection.

The performance hierarchy across models is clear (Protenix-v2 and IntelliFold v2 substantially outperform AlphaFold-Multimer, OpenFold3, and Boltz-2), but its origins are less certain. Several evaluated models disclose limited detail on their training data, and recent structure prediction methods increasingly augment their training sets with sequence-only interaction data or self-distilled predicted complexes [Abramson et al., 2024]. However, our temporal analysis finds no systematic decline in accuracy on newer patents, and differences across the evaluated training-cutoff splits are small relative to the performance differences between models (Appendix J). These findings provide no temporal evidence that training-data overlap drives the observed performance hierarchy, although incomplete training-data disclosure for some of these models prevents us from fully excluding overlap or disentangling improvements in binding prediction from differences in training-set coverage.

Our structural analyses show that a confident prediction is not necessarily a correct one: the best model contacts annotated epitope residues in 62.8% of cases, indicating that high ipSAE does not always imply a physically plausible binding mode. Cross-model structural consensus offers a complementary signal: when three or more models strongly agree on a pose, pairwise binder–decoy discrimination accuracy reaches 96.8%, but such agreement is rare, covering only 3% of pairs. We also find weak correlation between confidence and binding affinity, suggesting that these models capture a binary binding/non-binding distinction rather than a graded measure of interaction strength.

The experimental support behind POISK records varies with their patent sources. Some arise from focused characterisation studies reporting measured affinities against target homologues from multiple species or functional activity data; others arise from filings written mainly to protect sequence space. We recommend that users select records by the evidence needed for their application, for example filtering for reported binding affinity when required (Appendix C).

Beyond the benchmarking results, a key contribution of this work is the dataset itself. The patent literature is a largely untapped resource for computational biology, yet it contains one of the largest publicly available collections of experimentally validated antibody-antigen interactions. Our extraction pipeline yields over 89,000 records spanning more than 2000 unique targets, with rich per-record annotations including epitope residues, binding affinities, format, and discovery methodology. With only 716 entries shared with the recently published ANDD dataset, our collection is largely complementary to existing resources. We release the POISK dataset at https://zenodo.org/records/21775623, alongside an interactive webserver for data exploration at https://poisk-explorer.github.io/, to support future benchmarking, training data curation, and the development of antibody-specific scoring functions as new models continue to emerge.

## Appendix

### A Dataset fields

Table 3 presents the full list of the patent-extracted fields. The published POISK dataset also includes audit fields that provide additional information regarding the data accuracy for each specific record.

**Table 3:** List of extracted field names including audit fields. While the majority of these fields were extracted directly from explicit textual evidence, the LLM model was tasked with inferring two latent attributes: antibody_discovery_status (classifying sequences as novel inventions or referenced prior art) and uniprot_id (either directly mentioned or mapped from target names and organism data).

| Field Name |
|---|
| antibody_identifier |
| antibody_type |
| antibody_discovery_status |
| vh.seq_id |
| vh.is_full_heavy_chain |
| vl.seq_id |
| vl.is_full_light_chain |
| similar_antibody_seq_ids |
| discovery_methodology.cdr_modified |
| discovery_methodology.source_organism |
| discovery_methodology.specific_platform |
| discovery_methodology.strategies |
| target.organism |
| target.target_type |
| target.protein_name |
| target.gene_names |
| target.uniprot_id |
| target.seq_id |
| target.domain_start |
| target.domain_end |
| target.domain_coordinates |
| target.binding_components |
| target.binding_affinity.affinity_value |
| target.binding_affinity.method |
| target.binding_conditions |
| epitope.type |
| epitope.linear_epitope_range.start |
| epitope.linear_epitope_range.end |
| epitope.structural_contact_residues |
| epitope.antigenic_site_name |
| epitope.binning_competitors |
| epitope.evidence_types |
| epitope.functional_properties |
| epitope.ligand_receptor_competition |
| epitope.mutagenesis_residues |
| epitope.variants |
| audit.verdict |
| audit.flags |
| audit.reasoning |

Some patent documents described hundreds of antibody chains sharing a target (e.g., *“A heavy chain can be selected from SEQ ID NOs: 10–33, 38–80, and 160–183 and 222–245,”* patent US8679502). Gemini was instructed to pair and group these chains under a similar_antibody_seq_ids field.

### B Manual validation of the extraction pipeline

POISK rests on an LLM extraction step, and the automated Gemini auditor described in Section 2.1 validates records using the same model that was used to extract them. To obtain an independent estimate of extraction quality we therefore carried out a manual audit against the source patents.

#### Protocol

We drew two independent random samples of 20 records: one from the co-folding benchmark subset (11,247 records) and one from the full POISK dataset (89,074 records). A single experienced annotator retrieved each source patent and checked, field by field, against the patent text:

- correct *V_H_* /*V_L_*chain pairing, including whether a chain reported as absent is genuinely absent from the patent;
- correct antigen protein identity;
- correct antigen organism;
- correct binding-affinity value, for records that report one;
- correct epitope information, for records that report it (linear range, structural contact residues, antigenic site name, and mutagenesis residues).

A record was scored as fully correct only if every populated field passed. The 40-record audit required approximately six hours of annotator time, which is what sets the sample size: manual verification at this depth does not scale to a meaningful fraction of an 89,000-record resource.

#### Results

Table 4 reports per-field results. Denominators differ across fields because not every record reports every field; the denominator is the number of records in the sample for which the field is populated and therefore auditable.

**Table 4:**
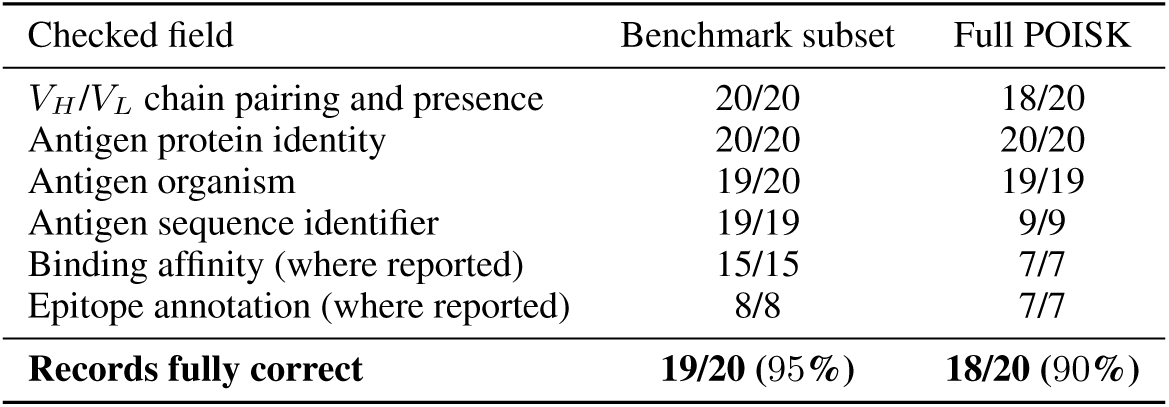
Manual audit of 40 records against their source patents. Each cell gives correct/auditable, where “auditable” counts the records in that sample of 20 for which the field is populated. Record-level accuracy requires every populated field of the record to be correct.

| Checked field | Benchmark subset | Full POISK |
|---|---|---|
| *V<sub>H</sub>*/*V<sub>L</sub>* chain pairing and presence | 20/20 | 18/20 |
| Antigen protein identity | 20/20 | 20/20 |
| Antigen organism | 19/20 | 19/19 |
| Antigen sequence identifier | 19/19 | 9/9 |
| Binding affinity (where reported) | 15/15 | 7/7 |
| Epitope annotation (where reported) | 8/8 | 7/7 |
| **Records fully correct** | **19/20** (95%) | **18/20** (90%) |

In the benchmark subset the single discrepancy was the target.organism field of antibody XENP31726 (patent US 11739145, record 76,346), labelled *Humanized* rather than *Human*. The target sequence itself was extracted correctly, so the folding benchmark is unaffected by this error.

In the full POISK sample both errors were auditor false-negatives on chain presence: the *V_L_*of record 4811 (US 7657380) and the *V_H_*of record 47,819 (US 10590187) were recorded as NOT_APPLICABLE, but the missing chain is in fact retrievable from the patent text. This is the expected direction of failure for an auditor configured to discard anything it cannot ground explicitly, and it is removed by construction from the benchmark subset, which requires both chains to be present. One further record in the full POISK sample (56,009, a non-targeting control antibody) legitimately carries no antigen organism and is excluded from that field’s denominator rather than counted as an error.

No spurious antibody–antigen binding relationships, that is, records asserting a binding pair that the patent does not support, were observed in either sample. We report the audit as evidence that the pipeline’s error modes are the ones we expected (conservative field dropping, occasional organism-label imprecision) rather than as a tight estimate of dataset-wide precision.

#### Audited record identifiers

Each record can be located in the released POISK dataset by its unique_record_id.

- **Benchmark subset (**11,247 **set):** 3987, 11225, 14896, 21130, 34392, 37911, 43201, 43930, 44635, 50741, 66444, 69183, 71114, 75364, 76346, 78250, 81637, 81741, 82552, 83896.
- **Full POISK (**89,074 **set):** 1686, 4811, 6294, 7782, 16977, 23850, 29531, 36301, 40645, 41452, 45559, 47819, 50822, 53537, 56009, 58606, 68268, 79907, 80406, 82111.

### C Evidential strength and evidence-tier filtering

Patent disclosures differ in how strongly they evidence a binding claim. Some records come from focused characterisation studies that report a measured affinity and a mapped epitope; others come from filings whose primary purpose is to protect sequence space around a lead. This heterogeneity is intrinsic to any patent-derived resource, and users of POISK should select an evidence tier appropriate to their application rather than treat all records as equivalent.

Two properties of the pipeline already raise the floor. The Gemini extraction prompt is instructed to emit an antibody–antigen pair only where the patent provides explicit evidence of binding, and the benchmark subset additionally requires a concrete antigen sequence, which removes claims that name no specific target. Beyond that floor, we recommend filtering on the annotation fields themselves.

Table 5 gives the size of each tier within the 11,247 benchmark binders. A reported binding affinity is the strongest single indicator that a record derives from experimental characterisation rather than from a sequence-space claim, and it is available for a little over half the benchmark.

**Table 5:**
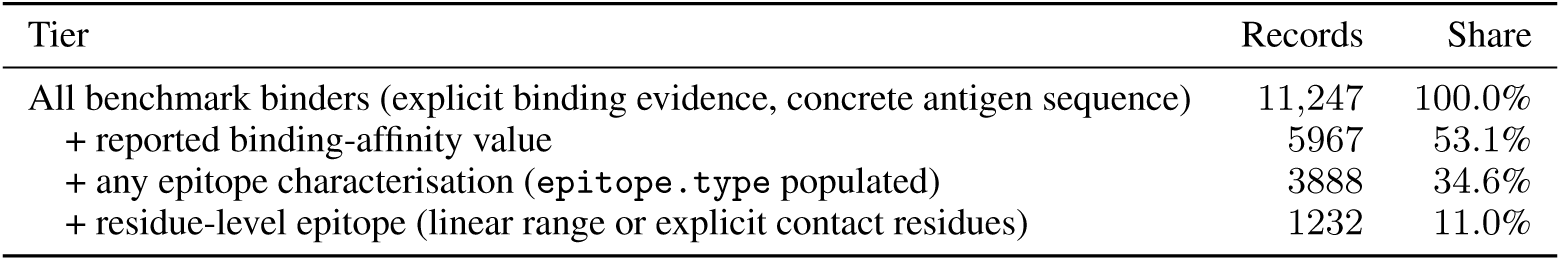
Evidence tiers among the 11,247 binders of the co-folding benchmark. These subsets overlap and do not represent successive filtering steps: a record may carry an affinity without an epitope, or vice versa.

Within the 3888 records carrying an epitope characterisation, 1003 are annotated as discontinuous or conformational, 826 as linear, 501 as functional or competitive, and 1558 as not further specified. Affinity values are stored as the free text reported in the patent (for example *2.30 nM* or *Kd=1.7×10^-9 M*); Section 3.2 standardises the 5207 of these that parse unambiguously into a numeric pAffinity, and Section 3.3 uses 1232 records for the epitope-recovery analysis.

Sequence-space filings are also not the dominant contributor to record count. POISK contains 78,174 unique antibody sequences across 10,110 patents, and the 11,247 benchmark binders come from 1546 distinct patents. A large per-patent sequence count moreover does not by itself imply weak per-record evidence: the largest contributor after removing duplicate sequences, US 11897968, discloses 87 unique antibody sequences and reports an SPR-measured binding affinity for each of them. The smaller filings that dominate the record count more typically correspond to focused characterisation studies.

### D Putative negatives: biological justification and label-noise sensitivity

The benchmark’s non-binders are putative: they are antibodies that we have strong reason to believe do not bind the paired antigen, not antibodies experimentally shown not to bind it. This appendix sets out why we expect high-affinity false negatives to be rare, and then bounds what residual mislabelling could do to the reported accuracies without relying on that expectation.

#### D.1 Why high-affinity false negatives should be rare

Of the 11,247 decoys, 7647 (68.0%) are retrieved from the Observed Antibody Space with KA-Search and 3600 (32.0%) are generated by germline reversion with CDR-H3 randomisation. Both classes are paired against human antigens.

For the KA-Search decoys, the source repertoires are those of healthy donors. High-affinity antibodies against human self-proteins are removed from the naive repertoire by central and peripheral tolerance (clonal deletion, receptor editing, anergy), and their systematic escape presents clinically as autoimmunity. The self-reactivity that does survive in healthy repertoires is dominated by polyreactive natural antibodies, which bind their many targets with low affinity, *K_D_* = 10*^−^*^3^ to 10*^−^*^7^ mol l*^−^*^1^, against 10*^−^*^7^ to 10*^−^*^11^ mol l*^−^*^1^ for specific monoreactive antibodies [Zhou et al., 2007]. The binders these decoys are paired against sit within that second range: of the 5967 benchmark binders that carry a reported affinity, 5207 (87.3%) give a value that parses to a molar concentration; their median is 1.0 nM, 80.2% are at or below 10 nM and 95.0% at or below 100 nM. (10.3% of the affinity strings are explicitly IC_50_ or EC_50_ rather than *K_D_*, so this is a distribution over reported affinity and potency values, not over *K_D_*alone.) Retrieving, by CDR similarity to an unrelated antibody, a healthy-donor sequence that happens to have specific complementarity to a given human self-protein at that strength is therefore unlikely.

For the germline-reverted decoys, reversion strips the somatic hypermutations selected during affinity maturation, which is why germline precursors of therapeutic antibodies typically bind their cognate target orders of magnitude more weakly than the mature molecule. Randomising CDR-H3 on top of reversion returns the sequence towards a naive combinatorial state. Naive human antibody libraries reach diversities of at least 3.5 *×* 10^10^ unique molecules [Glanville et al., 2009]; a specific antigen is bound by a vanishing fraction of that space, which is precisely why panning campaigns are needed to recover binders from it.

Neither argument is a measurement, and we do not claim a numerical false-negative rate. What we can say empirically is that the two decoy classes, which fail in different ways if the argument above is wrong, yield closely comparable discrimination performance for every engine (Figure 10). A large population of inadvertent binders in one class but not the other would be expected to show up as a divergence between the two curves, and it does not.

#### D.2 Label-noise sensitivity

Each benchmark datapoint is a triplet (target, binder, decoy) in which the decoy is derived from the binder. If a triplet is mislabelled, that is, if the binder does not in fact bind or the decoy does, the comparison degenerates into ranking two adversarially similar sequences that should carry the same label. Writing *A_obs_* for the observed pairwise accuracy, *A_true_* for accuracy on correctly labelled triplets, *A_mis_* for accuracy on mislabelled ones, and *f* for the mislabelled fraction:

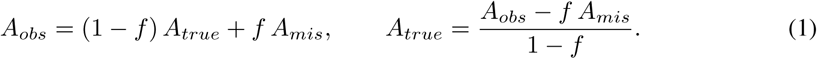

If a model has no systematic preference on degenerate triplets, then *A_mis_ ≈* 0.5, and since every engine we evaluate has *A_obs_ >* 0.5 it follows that *A_true_ > A_obs_* for any *f >* 0. Under that assumption label noise dilutes the reported accuracies rather than inflating them, and the numbers in Table 1 are conservative.

The assumption *A_mis_≈* 0.5 is the part that needs testing, since a model could plausibly prefer patent-derived sequences over repertoire-derived ones for reasons unrelated to the target. This assumption carries the whole argument and so we tested it directly.

##### Design

We sampled 300 benchmark datapoints under a target-uniqueness constraint, requiring distinct UniProt accessions and at most 50% pairwise sequence similarity between targets, and held decoy provenance balanced at 150 germline-reverted and 150 OAS-matched. We then shuffled the targets within the sampled set, so that in each resulting triplet neither the patent-derived sequence nor the decoy binds the antigen it is folded against; the target-uniqueness constraint is what guarantees that shuffling actually invalidates every triplet rather than occasionally reassigning a target to a genuine binder. The entire subset-sampling procedure was repeated under five sampling seeds (42–46), and every structure was folded under standard conditions with five prediction seeds and default parameters, using one high-accuracy engine (Protenix-v2) and one lower-accuracy engine (Boltz-2). This totals 30,000 structure predictions: 300 datapoints *×* 2 folds per datapoint *×* 5 sampling seeds *×* 2 engines *×* 5 predictions each.

We then measured how often the patent-derived sequence received the higher ipSAE. Pairs in which the engine assigns ipSAE exactly zero to both members carry no signal and are excluded; the “surviving pairs” column of Table 6 reports how many remain, and the accuracies are computed over those. The same sampled binder/decoy pairs were also scored against their original, correct targets using the archived benchmark folds, which checks that the sampled subsets are representative of overall benchmark performance rather than an unusually easy or hard slice.

**Table 6:**
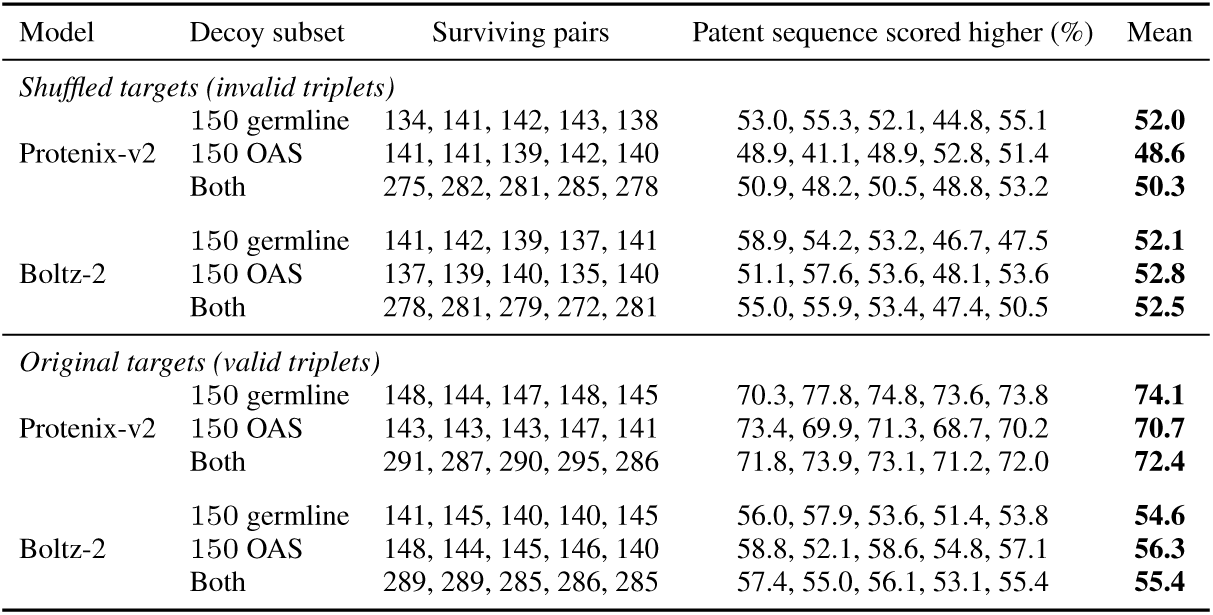
Discrimination on deliberately invalid (target-shuffled) triplets, and on the same pairs with their original correct targets. Per-seed values are listed in sampling-seed order 42, 43, 44, 45, 46. “Surviving pairs” excludes pairs for which the engine assigned an ipSAE of exactly zero to both the patent-derived sequence and the decoy; percentages are computed over surviving pairs. On the shuffled triplets, 50% indicates no systematic preference for the patent-derived sequence.

| Model | Decoy subset | Surviving pairs | Patent sequence scored higher (%) | Mean |
|---|---|---|---|---|
| *Shuffled targets (invalid triplets)* |  |  |  |  |
| Protenix-v2 | 150 germline | 134, 141, 142, 143, 138 | 53.0, 55.3, 52.1, 44.8, 55.1 | **52.0** |
|  | 150 OAS | 141, 141, 139, 142, 140 | 48.9, 41.1, 48.9, 52.8, 51.4 | **48.6** |
|  | Both | 275, 282, 281, 285, 278 | 50.9, 48.2, 50.5, 48.8, 53.2 | **50.3** |
| Boltz-2 | 150 germline | 141, 142, 139, 137, 141 | 58.9, 54.2, 53.2, 46.7, 47.5 | **52.1** |
|  | 150 OAS | 137, 139, 140, 135, 140 | 51.1, 57.6, 53.6, 48.1, 53.6 | **52.8** |
|  | Both | 278, 281, 279, 272, 281 | 55.0, 55.9, 53.4, 47.4, 50.5 | **52.5** |
| *Original targets (valid triplets)* |  |  |  |  |
| Protenix-v2 | 150 germline | 148, 144, 147, 148, 145 | 70.3, 77.8, 74.8, 73.6, 73.8 | **74.1** |
|  | 150 OAS | 143, 143, 143, 147, 141 | 73.4, 69.9, 71.3, 68.7, 70.2 | **70.7** |
|  | Both | 291, 287, 290, 295, 286 | 71.8, 73.9, 73.1, 71.2, 72.0 | **72.4** |
| Boltz-2 | 150 germline | 141, 145, 140, 140, 145 | 56.0, 57.9, 53.6, 51.4, 53.8 | **54.6** |
|  | 150 OAS | 148, 144, 145, 146, 140 | 58.8, 52.1, 58.6, 54.8, 57.1 | **56.3** |
|  | Both | 289, 289, 285, 286, 285 | 57.4, 55.0, 56.1, 53.1, 55.4 | **55.4** |

Both engines score invalid triplets at approximately 50% chance, for both decoy provenances, which is what the dilution argument requires. This holds for the strongest engine as well as the weakest: Protenix-v2 drops from 72.4% on valid pairs to 50.3% on invalid ones, Boltz-2 from 55.4 to 52.5%. Therefore, discrimination between binders and decoys is not explained by a systematic preference for patent-derived antibody sequences over generated negatives. Patent derived sequences scored better only when matched with the correct antigen.

This experiment required approximately 250 NVIDIA B200 GPU-hours, which is what limited it to 300 triplets and two engines.

### E Distribution of ipSAE scores

The density plots shown in Figure 8 illustrate population separation across the benchmarked folding models. The y-axis is truncated at a value of 20 to maintain visual clarity. This adjustment is necessary because a substantial volume of records receive an ipSAE score of exactly zero, which reflects the metric’s stringent confidence requirements for interface residues. Despite this significant clustering at zero, the ipSAE metric demonstrates the highest discriminatory power among all measured metrics. These results confirm that more recent architectures achieve a more distinct separation between binders and decoys than previous models.

**Figure 8:**
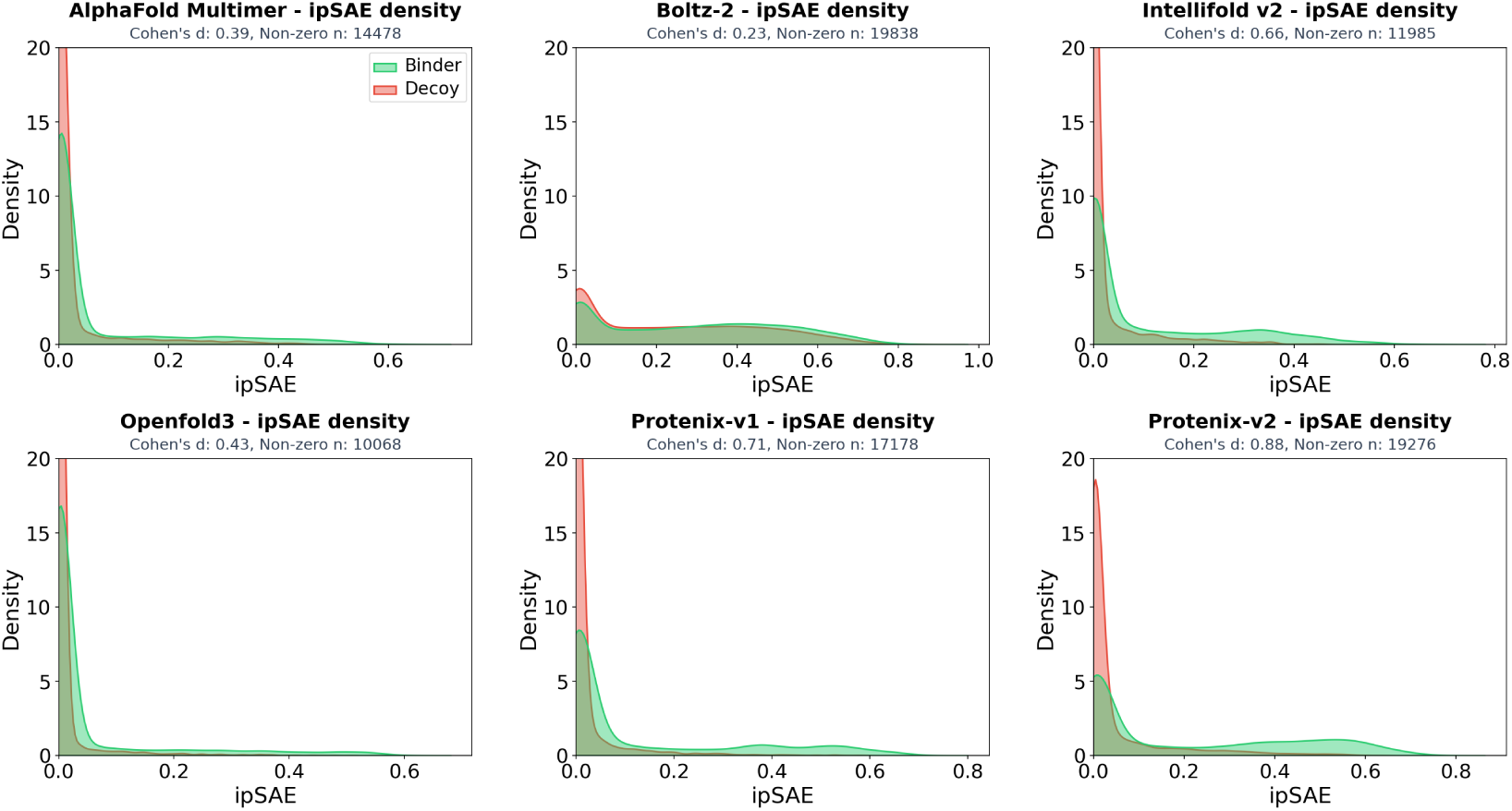
Baseline discrimination performance of six structure prediction models using the ipSAE metric. Kernel density estimates illustrate the separation between binder (green) and decoy (red) populations. The discriminatory power of each engine is quantified by Cohen’s *d*.

### F Pairwise accuracy as a function of confidence coverage

The headline pairwise accuracy quoted in Section 3.2.2 is measured after discarding the half of the pairs for which both members scored lowest, which mirrors how a practitioner would use a confidence filter but is not the accuracy over the benchmark as a whole. Table 7 reports the full trade-off: for each engine, pairwise accuracy at 100%, 75%, 50%, 25% and 10% coverage, where coverage is the fraction of the 11,247 pairs retained after ranking pairs by the higher of their two ipSAE scores and keeping the top fraction.

**Table 7:**
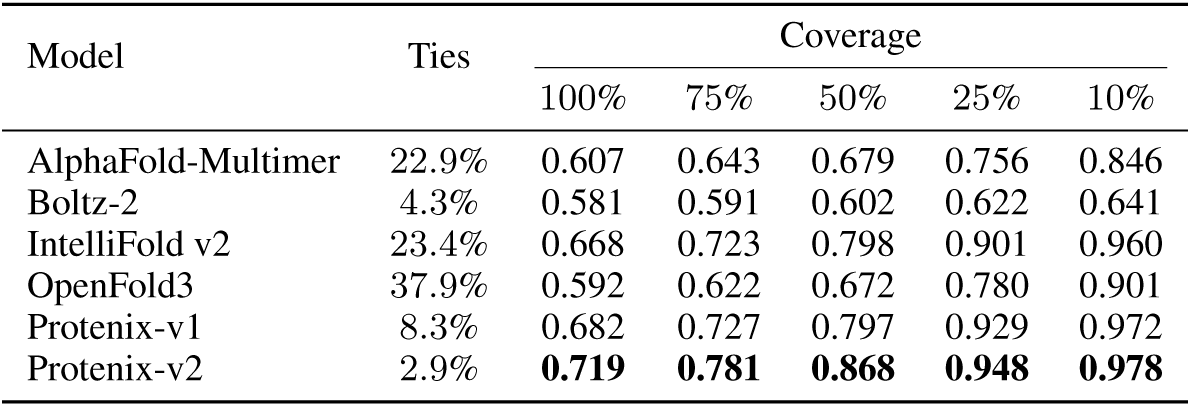
Pairwise binder-versus-decoy accuracy as a function of coverage. Coverage is the fraction of the 11,247 pairs retained, ranked by the maximum ipSAE within the pair. Exact ties receive half credit. The tie column is the fraction of all 11,247 pairs in which binder and decoy receive numerically identical ipSAE, which is what caps accuracy at full coverage.

| Model | Ties | Coverage |  |  |  |  |
|---|---|---|---|---|---|---|
|  |  | 100% | 75% | 50% | 25% | 10% |
| AlphaFold-Multimer | 22.9% | 0.607 | 0.643 | 0.679 | 0.756 | 0.846 |
| Boltz-2 | 4.3% | 0.581 | 0.591 | 0.602 | 0.622 | 0.641 |
| IntelliFold v2 | 23.4% | 0.668 | 0.723 | 0.798 | 0.901 | 0.960 |
| OpenFold3 | 37.9% | 0.592 | 0.622 | 0.672 | 0.780 | 0.901 |
| Protenix-v1 | 8.3% | 0.682 | 0.727 | 0.797 | 0.929 | 0.972 |
| Protenix-v2 | 2.9% | **0.719** | **0.781** | **0.868** | **0.948** | **0.978** |

Two features of this table are worth drawing out. First, accuracy increases monotonically as coverage tightens for every engine, which is the behaviour a usable confidence signal must show and is the substantive justification for reporting a filtered number at all. Second, the top and bottom of the ranking are stable across the whole range, so the conclusions of Section 3.2 do not depend on the chosen ipSAE cut-off: Protenix-v2 leads at every coverage level, and Boltz-2 trails at every coverage level.

The tie column accounts for part of the compression at full coverage. When both members of a pair fail to produce a confident interface, ipSAE returns exactly zero for both and the pair is undecidable; such pairs are scored at 0.5 here rather than dropped. OpenFold3 leaves 37.9% of pairs undecidable in this way and Protenix-v2 only 2.9%, so a substantial part of the gap between them at 100% coverage reflects how often each engine produces any interface prediction at all, not how well it ranks the ones it produces. This motivates reporting both full-benchmark accuracy and accuracy on the most confident half of pairs.

### G Benchmark stability across decoy strategies and antigen lengths

The robustness of the ipSAE metric is further characterized by evaluating its performance across distinct negative control groups and target size distributions. While the main results establish a baseline for binding classification, Figure 10 demonstrates that accuracy remains consistent regardless of whether non-binders are identified via sequence-based search (OAS) or generated through germline reversion with CDR-H3 randomisation. Additionally, the relationship between target length and model confidence is detailed in Figure 9. The trend of decreasing antigen length at higher confidence thresholds is visible for most models, contrasting with the anomalous positive correlation found in the IntelliFold v2 predictions.

**Figure 9:**
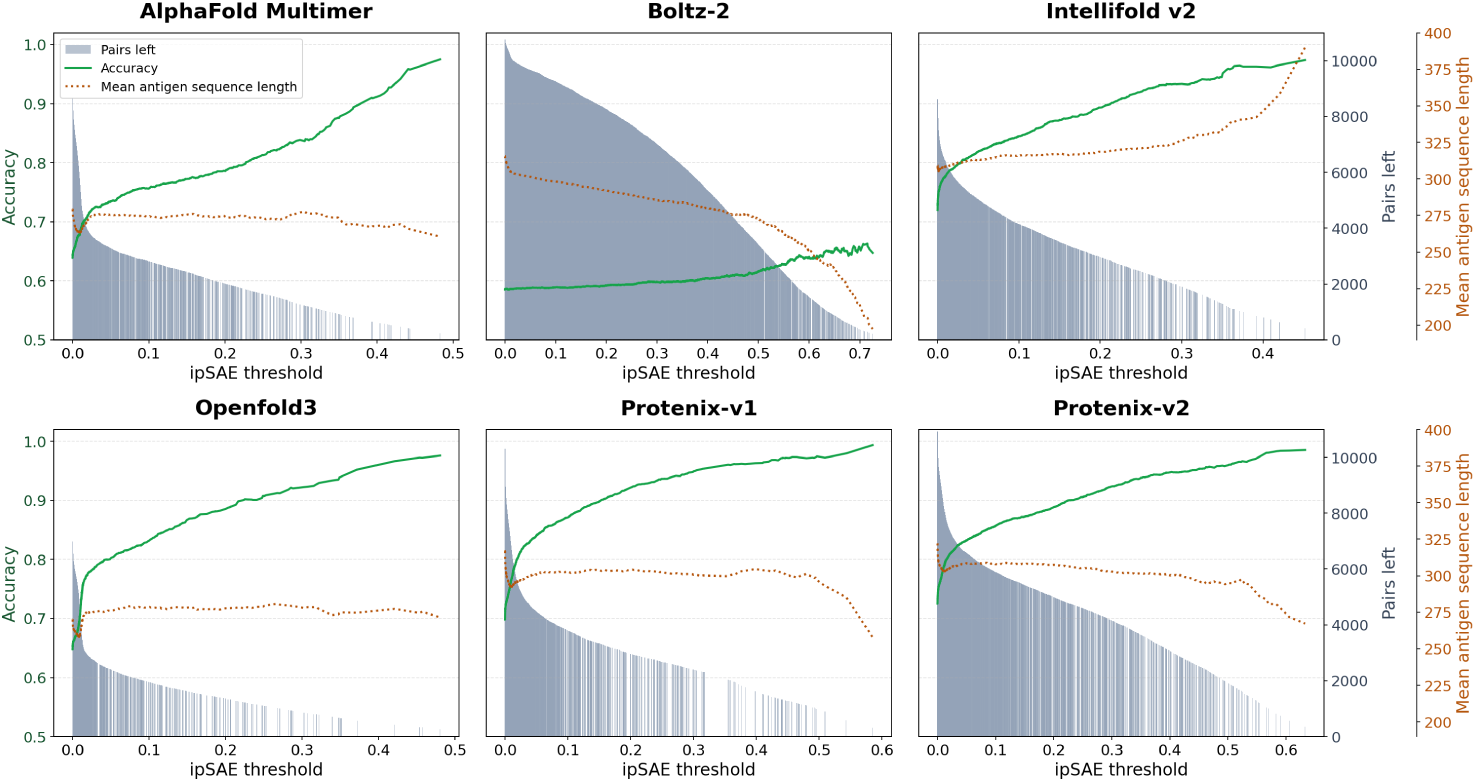
Relationship between target sequence length and structural confidence. Mean antigen sequence length (dashed line) is tracked as the ipSAE filtering threshold increases. While the average target length remains stable or decreases for most engines at higher confidence levels, IntelliFold v2 shows a positive correlation between confidence and target size.

**Figure 10:**
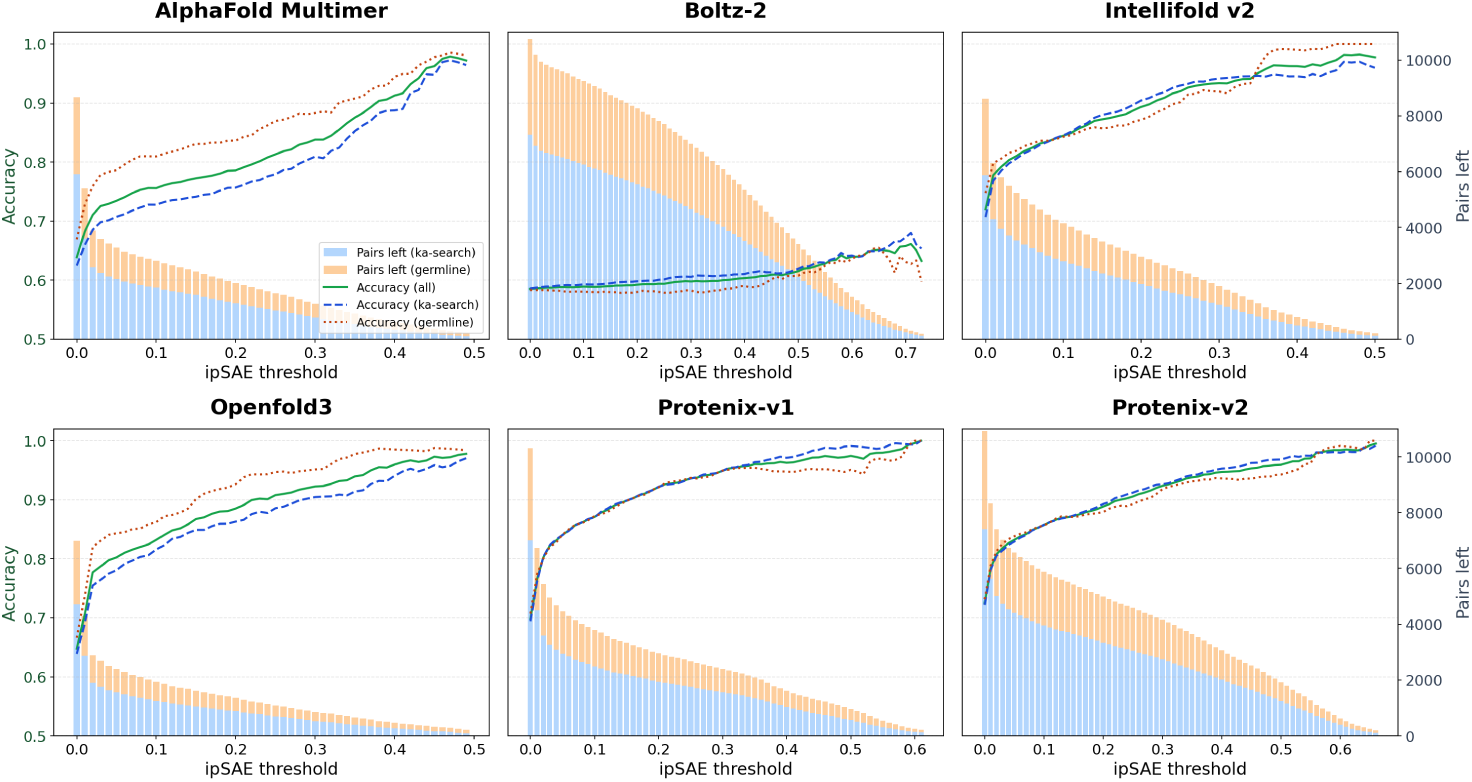
Model accuracy across different decoy generation strategies. Predictive performance is compared for complex pairs involving OAS KA-Search derived non-binding decoys versus those generated through germline reversion. Accuracy curves and pair retention counts indicate that the discriminatory power of the ipSAE metric is robust across different types of challenging decoys.

### H Target-dependent performance in the 1-in-50 screening simulation

The aggregate screening results summarise performance across 27 antigen constructs. To examine how this performance varies between targets, we report top-1 accuracy separately for each construct (Table 8). Each value averages results from 25 sampling seeds, with 20 trials per construct per seed, using the same runs as Figure 5.

**Table 8:**
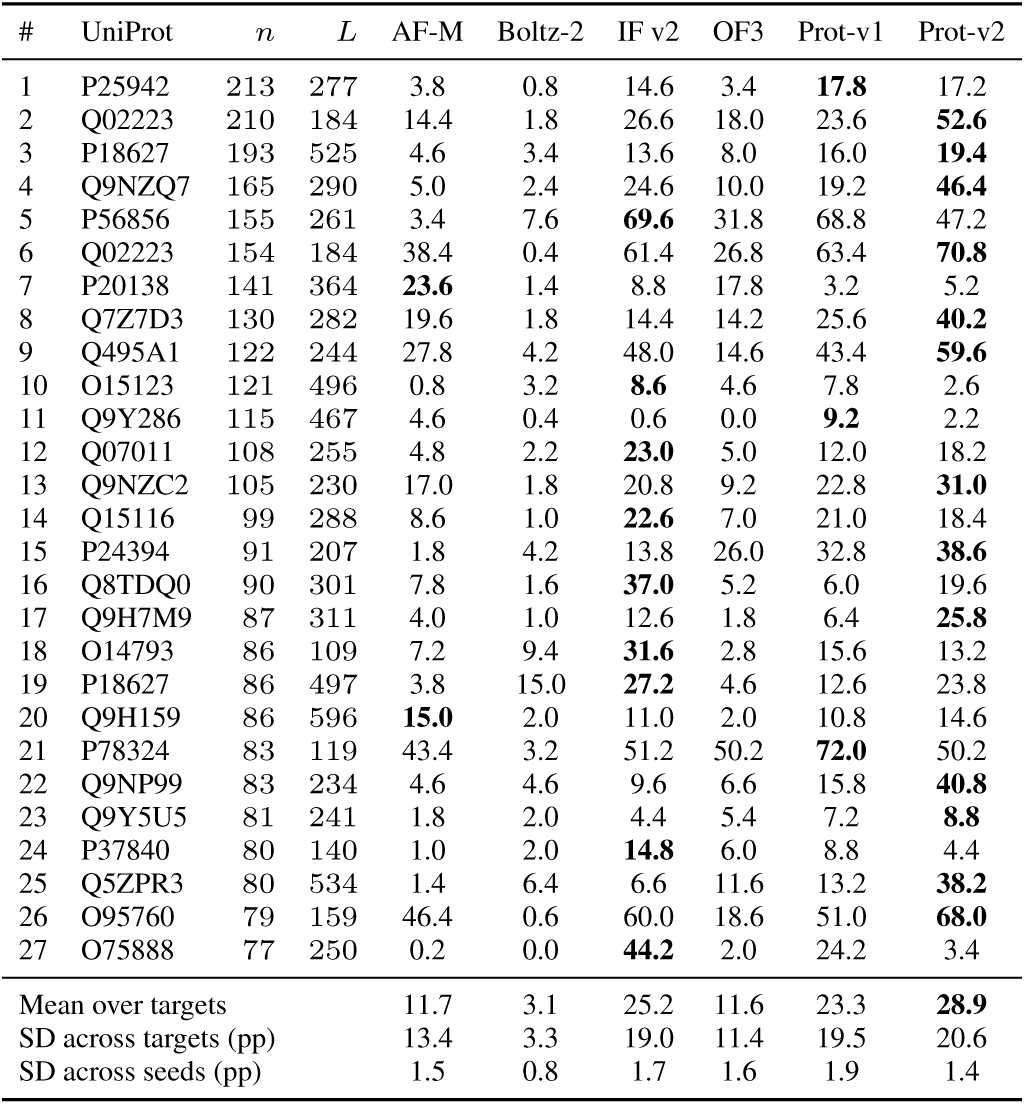
Top-1 accuracy (%) in the 1-in-50 screening simulation for each antigen construct. Each cell reports the mean over 25 sampling seeds with 20 trials per construct per seed. Top-1 success requires the binder to score strictly above all 49 decoys. *n* denotes the number of available patent-derived binders and *L* the antigen construct length in residues. Distinct constructs can share a UniProt accession. Bold values indicate the highest accuracy in each row; the random-selection baseline is 2%. Summary rows report the mean accuracy across targets (%) and standard deviations across targets and sampling seeds (percentage points).

| # | UniProt | *n* | *L* | AF-M | Boltz-2 | IF v2 | OF3 | Prot-v1 | Prot-v2 |
|---|---|---|---|---|---|---|---|---|---|
| 1 | P25942 | 213 | 277 | 3.8 | 0.8 | 14.6 | 3.4 | **17.8** | 17.2 |
| 2 | Q02223 | 210 | 184 | 14.4 | 1.8 | 26.6 | 18.0 | 23.6 | **52.6** |
| 3 | P18627 | 193 | 525 | 4.6 | 3.4 | 13.6 | 8.0 | 16.0 | **19.4** |
| 4 | Q9NZQ7 | 165 | 290 | 5.0 | 2.4 | 24.6 | 10.0 | 19.2 | **46.4** |
| 5 | P56856 | 155 | 261 | 3.4 | 7.6 | **69.6** | 31.8 | 68.8 | 47.2 |
| 6 | Q02223 | 154 | 184 | 38.4 | 0.4 | 61.4 | 26.8 | 63.4 | **70.8** |
| 7 | P20138 | 141 | 364 | **23.6** | 1.4 | 8.8 | 17.8 | 3.2 | 5.2 |
| 8 | Q7Z7D3 | 130 | 282 | 19.6 | 1.8 | 14.4 | 14.2 | 25.6 | **40.2** |
| 9 | Q495A1 | 122 | 244 | 27.8 | 4.2 | 48.0 | 14.6 | 43.4 | **59.6** |
| 10 | O15123 | 121 | 496 | 0.8 | 3.2 | **8.6** | 4.6 | 7.8 | 2.6 |
| 11 | Q9Y286 | 115 | 467 | 4.6 | 0.4 | 0.6 | 0.0 | **9.2** | 2.2 |
| 12 | Q07011 | 108 | 255 | 4.8 | 2.2 | **23.0** | 5.0 | 12.0 | 18.2 |
| 13 | Q9NZC2 | 105 | 230 | 17.0 | 1.8 | 20.8 | 9.2 | 22.8 | **31.0** |
| 14 | Q15116 | 99 | 288 | 8.6 | 1.0 | **22.6** | 7.0 | 21.0 | 18.4 |
| 15 | P24394 | 91 | 207 | 1.8 | 4.2 | 13.8 | 26.0 | 32.8 | **38.6** |
| 16 | Q8TDQ0 | 90 | 301 | 7.8 | 1.6 | **37.0** | 5.2 | 6.0 | 19.6 |
| 17 | Q9H7M9 | 87 | 311 | 4.0 | 1.0 | 12.6 | 1.8 | 6.4 | **25.8** |
| 18 | O14793 | 86 | 109 | 7.2 | 9.4 | **31.6** | 2.8 | 15.6 | 13.2 |
| 19 | P18627 | 86 | 497 | 3.8 | 15.0 | **27.2** | 4.6 | 12.6 | 23.8 |
| 20 | Q9H159 | 86 | 596 | **15.0** | 2.0 | 11.0 | 2.0 | 10.8 | 14.6 |
| 21 | P78324 | 83 | 119 | 43.4 | 3.2 | 51.2 | 50.2 | **72.0** | 50.2 |
| 22 | Q9NP99 | 83 | 234 | 4.6 | 4.6 | 9.6 | 6.6 | 15.8 | **40.8** |
| 23 | Q9Y5U5 | 81 | 241 | 1.8 | 2.0 | 4.4 | 5.4 | 7.2 | **8.8** |
| 24 | P37840 | 80 | 140 | 1.0 | 2.0 | **14.8** | 6.0 | 8.8 | 4.4 |
| 25 | Q5ZPR3 | 80 | 534 | 1.4 | 6.4 | 6.6 | 11.6 | 13.2 | **38.2** |
| 26 | O95760 | 79 | 159 | 46.4 | 0.6 | 60.0 | 18.6 | 51.0 | **68.0** |
| 27 | O75888 | 77 | 250 | 0.2 | 0.0 | **44.2** | 2.0 | 24.2 | 3.4 |
| Mean over targets |  |  |  | 11.7 | 3.1 | 25.2 | 11.6 | 23.3 | **28.9** |
| SD across targets (pp) |  |  |  | 13.4 | 3.3 | 19.0 | 11.4 | 19.5 | 20.6 |
| SD across seeds (pp) |  |  |  | 1.5 | 0.8 | 1.7 | 1.6 | 1.9 | 1.4 |

Screening success varies substantially between targets. For Protenix-v2, top-1 accuracy ranges from 2.2% to 70.8%, with a standard deviation of 20.6 percentage points across targets, compared with 1.4 points across sampling seeds. Between-target variation exceeds sampling variation for every engine, indicating that the aggregate results reflect a broad range of screening difficulty. Three targets remain challenging for all six engines, with none exceeding 10% top-1 accuracy, whereas several engines perform well on other targets.

The best-performing engine also depends on the target. Protenix-v2 achieves the highest top-1 accuracy on 13 of the 27 constructs and IntelliFold v2 on 9, while Protenix-v1 and AlphaFold-Multimer lead on the remaining five. Thus, Protenix-v2’s advantage in the aggregate does not translate into uniformly superior performance on every antigen. These results complement the overall enrichment reported in the main text by showing how both screening difficulty and relative model performance depend on the target.

### I Structural confidence in relation to binding affinity and sequence novelty

Structural confidence could reflect not only whether an antibody binds, but also how strongly it binds. We examined this possibility by comparing ipSAE across intervals of reported affinity and potency (Figure 11). Higher-affinity records show a weak tendency towards higher confidence, but the broad overlap between score distributions limits the usefulness of ipSAE as a measure of binding strength.

**Figure 11:**
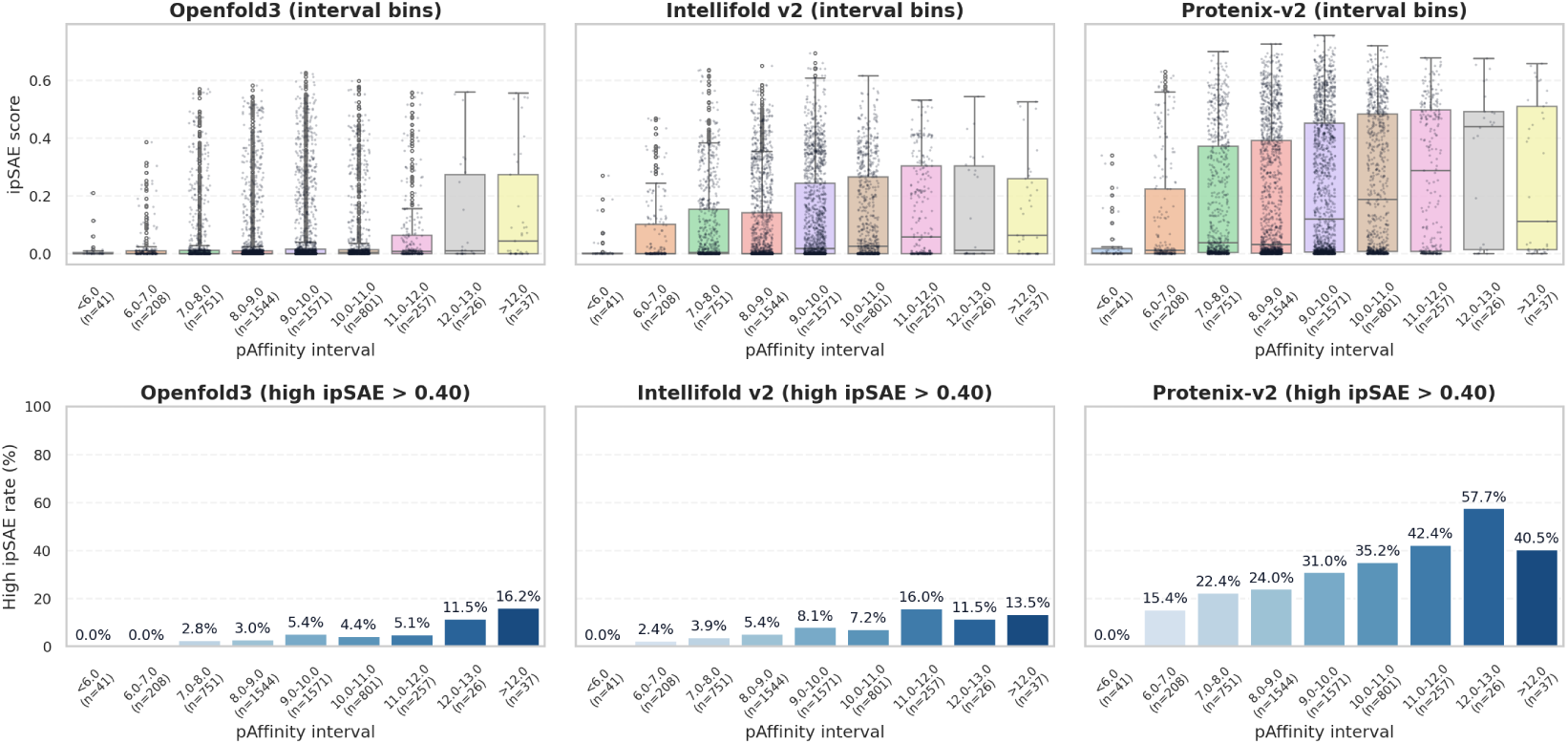
Relationship between binding affinity and structural confidence. The upper panels display the distribution of ipSAE scores across different pAffinity intervals for representative engines: OpenFold3, IntelliFold v2, and Protenix-v2. The lower panels plot the percentage of records in each affinity bin that achieve a high ipSAE score above 0.40. These results show a marginal trend where high-affinity records are more frequently associated with higher structural confidence.

We also examined whether confidence depends on similarity to antibodies with resolved structures. For binders targeting antigens with matching SAbDab entries, we compared ipSAE with the minimum cumulative CDR edit distance to antibodies against those targets (Figure 12). Confidence shows a weak tendency to decrease with increasing sequence distance, suggesting some preference for antibodies closer to known structures. However, this relationship is not strong enough for sequence proximity alone to explain the variation in confidence.

**Figure 12:**
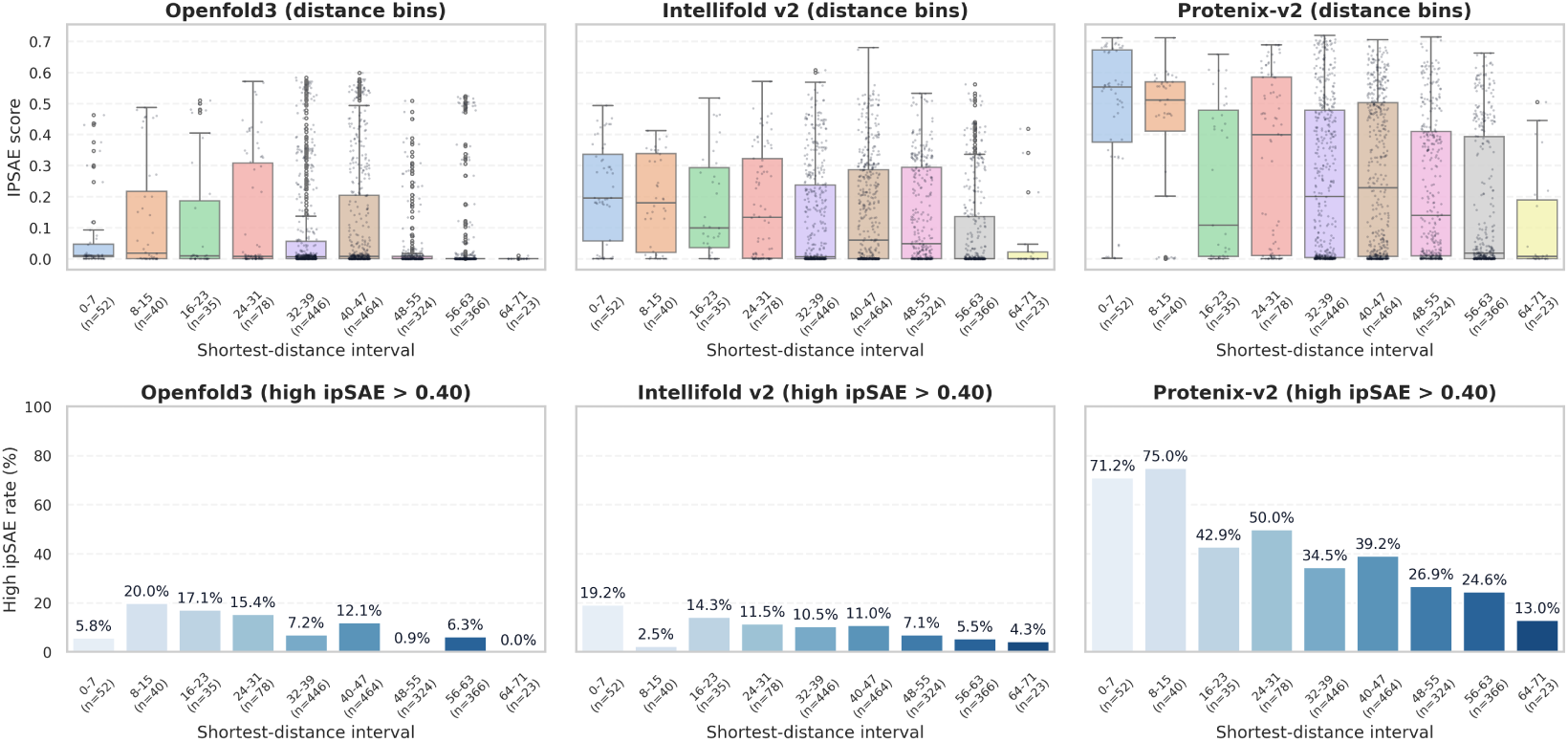
Impact of structural novelty on folding confidence. The analysis compares the shortest sequence distance to known SAbDab structures against predicted ipSAE scores. Upper panels display binned score distributions, and lower panels indicate the rate of high ipSAE scores per distance interval. High variance across intervals suggests that sequence novelty has a limited impact on model confidence.

### J Temporal analysis of potential training-data contamination

Several of the evaluated engines disclose little about their training corpora, so contamination by patent-derived text or sequences cannot be excluded by inspection. It can, however, be probed from the outside. Patent content becomes available for ingestion at a known time, and older patents have had longer to reach any corpus assembled before a model was trained. Systematic memorisation of patent content should therefore appear as higher accuracy on older records and a decline on the most recent ones.

We grouped every benchmark pair by the publication date of the binder’s source patent, using one bucket for pre-2018 patents and six-month buckets thereafter, running through H1 2025. The endpoint reflects the ingestion lag of our own pipeline rather than the patent record: sequence listings from a granted patent typically take between a few months and a year to appear in GenBank, and our extraction was run in January 2026. Each pair is scored with the paper’s primary ipSAE metric, with half credit for exact ties so that engines with high folding-failure rates are not penalised asymmetrically.

Accuracy does not systematically decline as patent publication dates become more recent (Figure 13). Across all six engines, performance on H1 2025 records is comparable to or better than on pre-2018 records. Protenix-v2 shows a small decrease, from 0.716 to 0.689, whereas IntelliFold v2 shows a small increase, from 0.654 to 0.669. Both changes are modest relative to the fluctuations across publication periods. The clearest shared pattern is higher accuracy on H2 2018 records: five of the six engines perform best in this period, while OpenFold3 peaks in H1 2019. This common pattern suggests that some cohorts contain interactions that are easier for models to distinguish, rather than showing a consistent advantage for older records. Taken together, these results provide no temporal evidence that preferential performance on older patents explains the differences between models.

**Figure 13:**
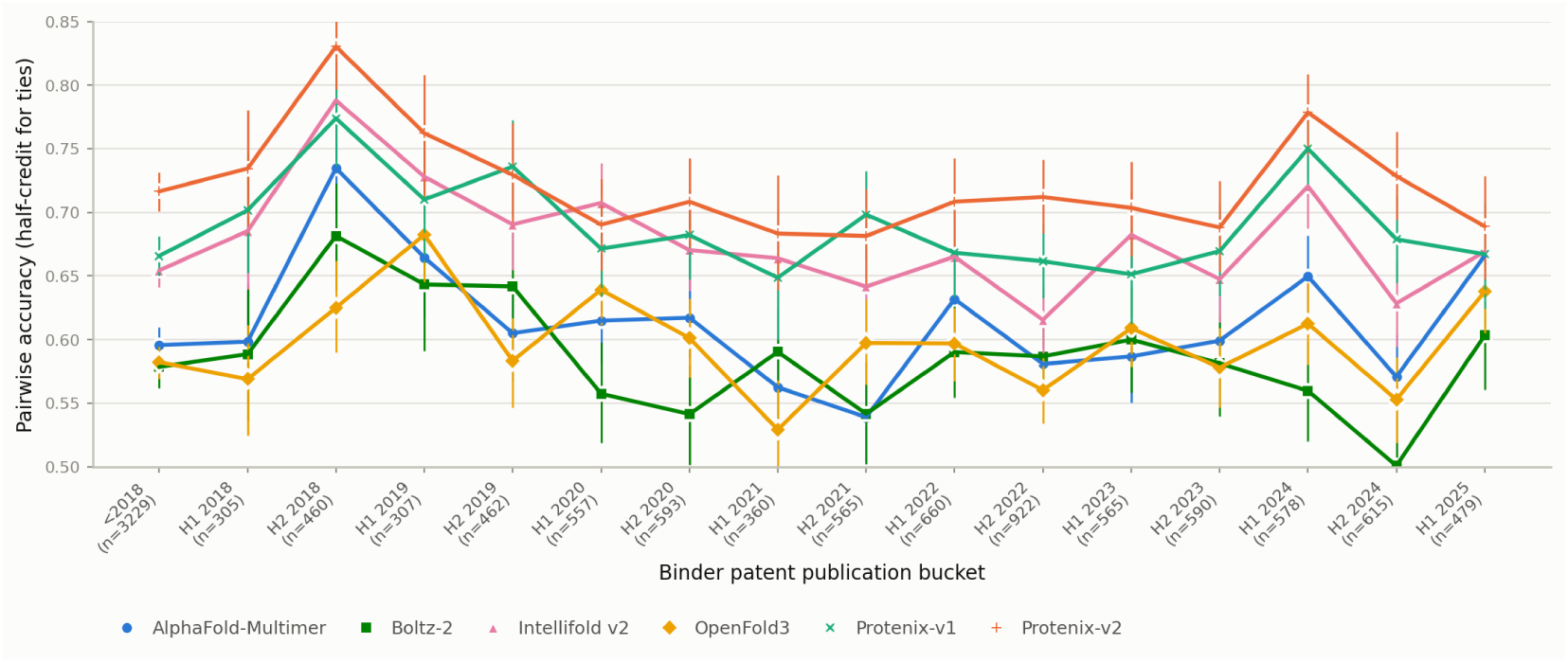
Per-bucket pairwise accuracy (half-credit for ties) across binder-patent publication buckets. One <2018 bucket collects pre-2018 pairs; from 2018 onwards, buckets are 6-month H1/H2 windows. Bucket sizes are given on the axis and range from 305 to 3229 pairs. Error bars are bootstrap 95% confidence intervals.

As a second, coarser test we split the benchmark at each engine’s own stated PDB training cutoff (Figure 14). Under memorisation we would expect accuracy to be higher before the cutoff than after it.

**Figure 14:**
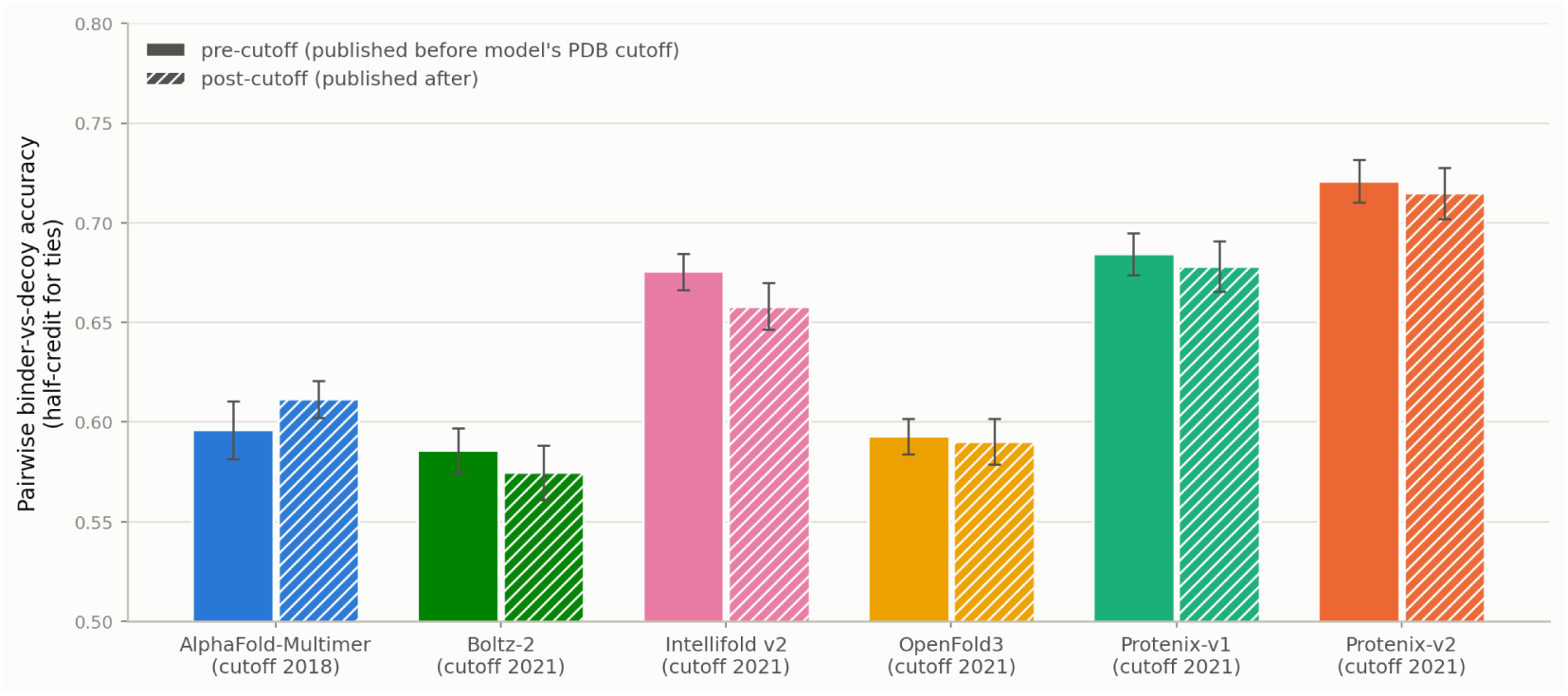
Pre-versus post-cutoff pairwise binder-versus-decoy accuracy. Solid bars are pairs whose patent published before each engine’s stated PDB training cutoff, hatched bars those published after. Pairs are scored with half-credit for ties. Error bars are bootstrap 95% confidence intervals. Pre-cutoff *n* is 3413 for AlphaFold-Multimer and 6616 for the other five engines, post-cutoff 7834 and 4631 respectively.

The largest pre-minus-post difference across all six engines is 1.8 percentage points (IntelliFold v2). For the two strongest engines, Protenix-v1 and Protenix-v2, the differences are 0.6 points. Every difference is under two points, against a spread between engines of more than ten points, so none of them is large enough to account for model ranking.

### K Structural consensus

High folding confidence does not necessarily imply structural agreement across models. To illustrate this distinction, we show predictions for Humanized ICR-1.1 from patent US5837822, a selected example in which confident predictions correspond to markedly different binding poses (Figure 15). Although the predicted antigen conformations are relatively consistent across the six evaluated folding engines, the models place the antibody at widely different epitopes. Their disagreement therefore extends beyond confidence-score calibration to the predicted geometry of the antibody-antigen interaction.

**Figure 15:**
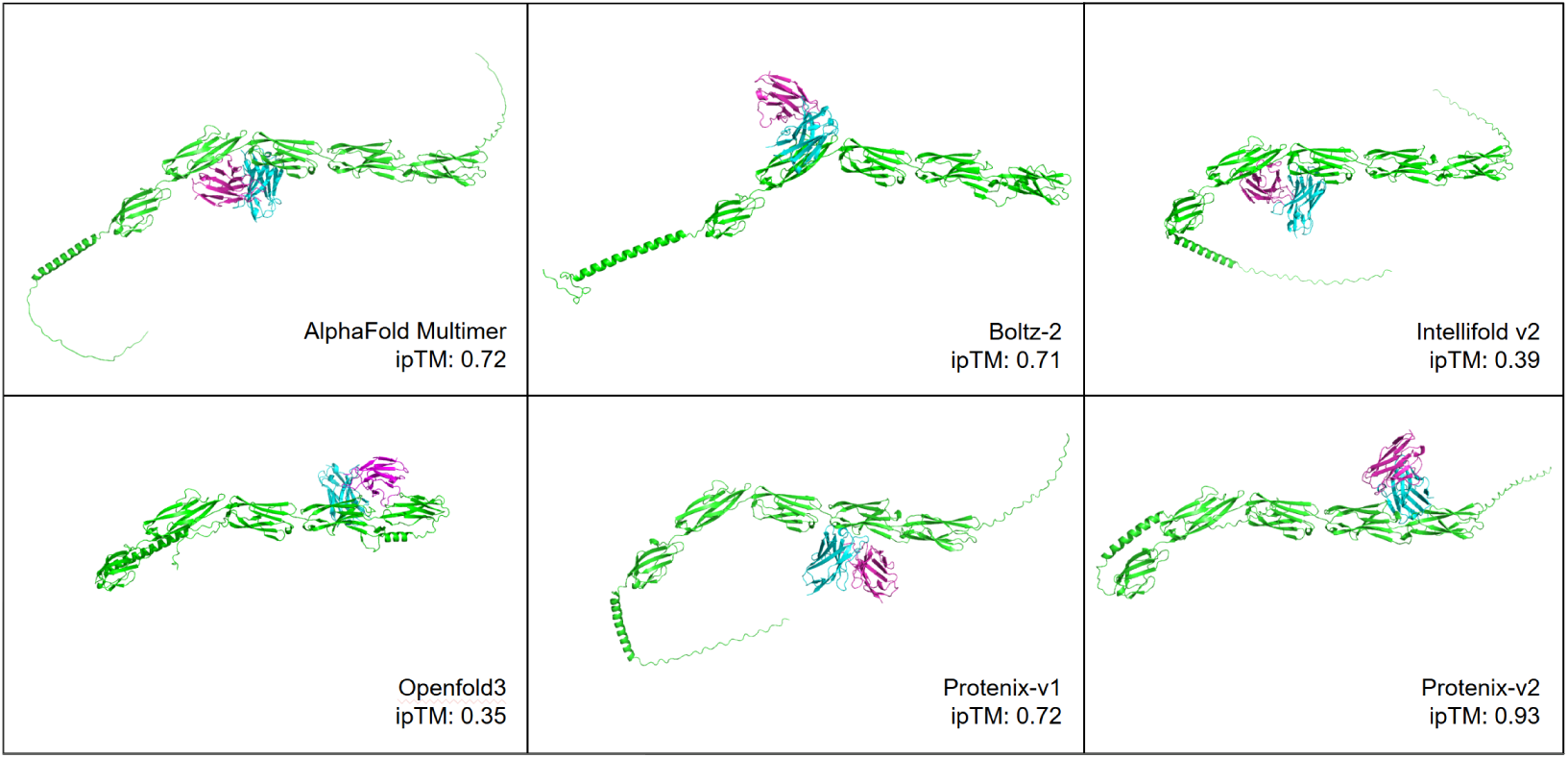
Structure predictions for Humanized ICR-1.1 from the patent US5837822.

Requiring agreement among five or six models further restricts coverage, extending the analysis in Section 3.3.1 (Figure 16). When all six models must agree (*k* = 6), approximately half of the dataset is excluded even at a DockQ threshold of 0.01. The retained pairs nevertheless show high binder–decoy discrimination accuracy: at *DockQ ≥* 0.23, accuracy reaches at least 90%. Thus, agreement across all six models can provide a strong binding signal even without close agreement on the precise pose, but at the cost of reduced coverage.

**Figure 16:**
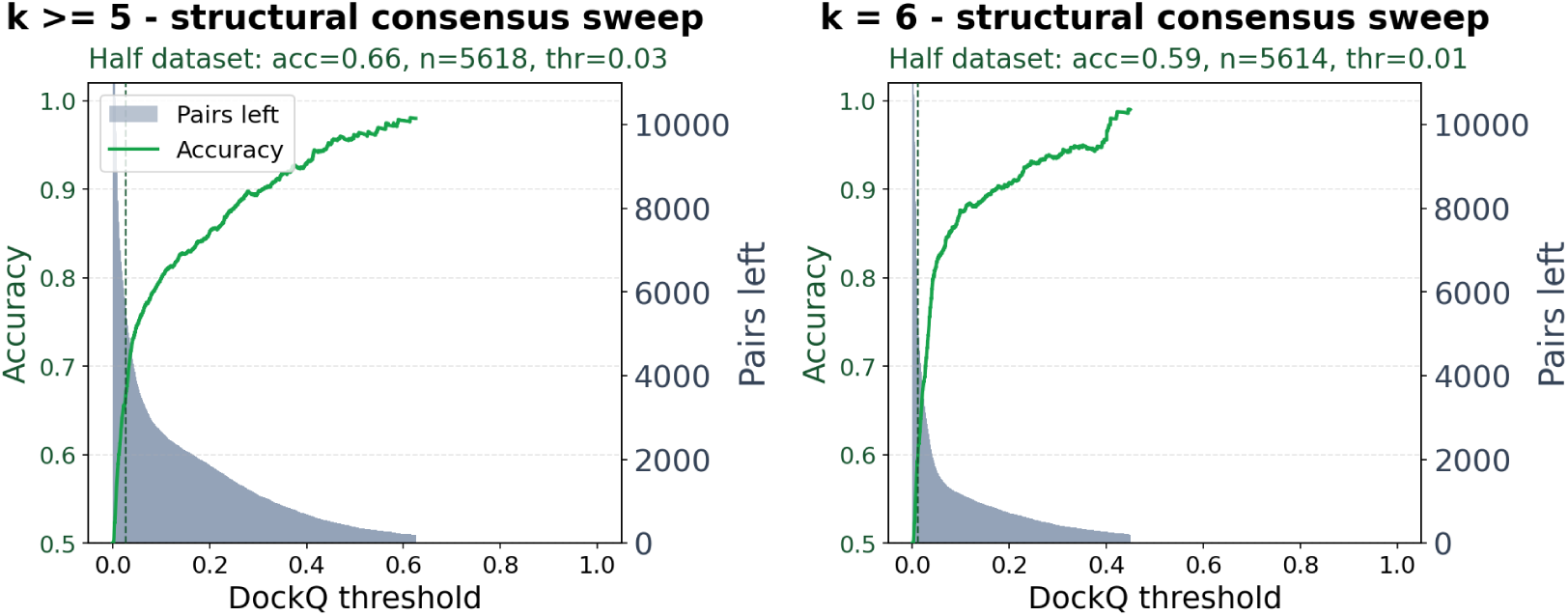
High-stringency structural-consensus binder/decoy filtering for *k ≥* 5 and *k* = 6. Bars show retained pairs and the green line shows pairwise accuracy as the DockQ threshold is tightened.

### L Compute and runtime

All folding was run on NVIDIA B200 GPUs. Each of the 22,494 complexes was folded by all six engines with five predictions and no templates, giving 674,820 structure predictions in total. MSAs were computed once per unique sequence with ColabFold and shared across engines and seeds, and are excluded from the reported timings.

**Table 9:** Total compute and per-model breakdown for benchmark refolding.

| Engine | B200 GPU-h |
|---|---|
| AlphaFold-Multimer | 2909 |
| Boltz-2 | 379 |
| IntelliFold v2 | 4027 |
| OpenFold3 | 535 |
| Protenix-v1 | 540 |
| Protenix-v2 | 641 |
| Total | 9031 |

#### NeurIPS Paper Checklist

##### 1. Claims

Question: Do the main claims made in the abstract and introduction accurately reflect the paper’s contributions and scope?

Answer: [Yes]

Justification: The abstract and introduction state three contributions—a large-scale patent-derived dataset, a benchmark of six folding models as zero-shot binding classifiers, and an analysis of structural consensus and epitope localization—all of which are substantiated in Sections 2–4 with quantitative results and appropriate caveats about limitations (e.g., putative negatives, possible data leakage).

Guidelines:

- The answer [N/A] means that the abstract and introduction do not include the claims made in the paper.
- The abstract and/or introduction should clearly state the claims made, including the contributions made in the paper and important assumptions and limitations. A [No] or [N/A] answer to this question will not be perceived well by the reviewers.
- The claims made should match theoretical and experimental results, and reflect how much the results can be expected to generalize to other settings.
- It is fine to include aspirational goals as motivation as long as it is clear that these goals are not attained by the paper.

##### 2. Limitations

Question: Does the paper discuss the limitations of the work performed by the authors?

Answer: [Yes]

Justification: The Discussion (Section 4) explicitly addresses several limitations: non-binding labels are putative and not experimentally validated, possible data leakage through undisclosed training data of evaluated models, weak correlation between confidence and affinity, poor epitope localization, and the inability to disentangle improvements in binding physics from training-set coverage differences.

Guidelines:

- The answer [N/A] means that the paper has no limitation while the answer [No] means that the paper has limitations, but those are not discussed in the paper.
- The authors are encouraged to create a separate “Limitations” section in their paper.
- The paper should point out any strong assumptions and how robust the results are to violations of these assumptions (e.g., independence assumptions, noiseless settings, model well-specification, asymptotic approximations only holding locally). The authors should reflect on how these assumptions might be violated in practice and what the implications would be.
- The authors should reflect on the scope of the claims made, e.g., if the approach was only tested on a few datasets or with a few runs. In general, empirical results often depend on implicit assumptions, which should be articulated.
- The authors should reflect on the factors that influence the performance of the approach. For example, a facial recognition algorithm may perform poorly when image resolution is low or images are taken in low lighting. Or a speech-to-text system might not be used reliably to provide closed captions for online lectures because it fails to handle technical jargon.
- The authors should discuss the computational efficiency of the proposed algorithms and how they scale with dataset size.
- If applicable, the authors should discuss possible limitations of their approach to address problems of privacy and fairness.
- While the authors might fear that complete honesty about limitations might be used by reviewers as grounds for rejection, a worse outcome might be that reviewers discover limitations that aren’t acknowledged in the paper. The authors should use their best judgment and recognize that individual actions in favor of transparency play an important role in developing norms that preserve the integrity of the community. Reviewers will be specifically instructed to not penalize honesty concerning limitations.

##### 3. Theory assumptions and proofs

Question: For each theoretical result, does the paper provide the full set of assumptions and a complete (and correct) proof?

Answer: [N/A]

Justification: The paper is an empirical benchmarking study and does not include theoretical results or proofs.

Guidelines:

- The answer [N/A] means that the paper does not include theoretical results.
- All the theorems, formulas, and proofs in the paper should be numbered and cross-referenced.
- All assumptions should be clearly stated or referenced in the statement of any theorems.
- The proofs can either appear in the main paper or the supplemental material, but if they appear in the supplemental material, the authors are encouraged to provide a short proof sketch to provide intuition.
- Inversely, any informal proof provided in the core of the paper should be complemented by formal proofs provided in appendix or supplemental material.
- Theorems and Lemmas that the proof relies upon should be properly referenced.

##### 4. Experimental result reproducibility

Question: Does the paper fully disclose all the information needed to reproduce the main experimental results of the paper to the extent that it affects the main claims and/or conclusions of the paper (regardless of whether the code and data are provided or not)?

Answer: [Yes]

Justification: Section 2 details the data extraction pipeline, filtering criteria, decoy construction strategy, folding protocol (MSA-based, no templates, five trunk calls with one diffusion sample), and scoring methodology (ipSAE with specified cutoffs). The dataset is released publicly on Zenodo (Section 3.1). All six evaluated models are publicly available or accessible.

Guidelines:

- The answer [N/A] means that the paper does not include experiments.
- If the paper includes experiments, a [No] answer to this question will not be perceived well by the reviewers: Making the paper reproducible is important, regardless of whether the code and data are provided or not.
- If the contribution is a dataset and/or model, the authors should describe the steps taken to make their results reproducible or verifiable.
- Depending on the contribution, reproducibility can be accomplished in various ways. For example, if the contribution is a novel architecture, describing the architecture fully might suffice, or if the contribution is a specific model and empirical evaluation, it may be necessary to either make it possible for others to replicate the model with the same dataset, or provide access to the model. In general. releasing code and data is often one good way to accomplish this, but reproducibility can also be provided via detailed instructions for how to replicate the results, access to a hosted model (e.g., in the case of a large language model), releasing of a model checkpoint, or other means that are appropriate to the research performed.
- While NeurIPS does not require releasing code, the conference does require all submissions to provide some reasonable avenue for reproducibility, which may depend on the nature of the contribution. For example
  (a) If the contribution is primarily a new algorithm, the paper should make it clear how to reproduce that algorithm.
  (b) If the contribution is primarily a new model architecture, the paper should describe the architecture clearly and fully.
  (c) If the contribution is a new model (e.g., a large language model), then there should either be a way to access this model for reproducing the results or a way to reproduce the model (e.g., with an open-source dataset or instructions for how to construct the dataset).
  (d) We recognize that reproducibility may be tricky in some cases, in which case authors are welcome to describe the particular way they provide for reproducibility. In the case of closed-source models, it may be that access to the model is limited in some way (e.g., to registered users), but it should be possible for other researchers to have some path to reproducing or verifying the results.

##### 5. Open access to data and code

Question: Does the paper provide open access to the data and code, with sufficient instructions to faithfully reproduce the main experimental results, as described in supplemental material?

Answer: [Yes]

Justification: The full patent-extracted dataset is released publicly on Zenodo (Section 3.1). The ipSAE scoring tool used is open source and linked in section 2.3.

Guidelines:

- The answer [N/A] means that paper does not include experiments requiring code.
- Please see the NeurIPS code and data submission guidelines (https://neurips.cc/public/guides/CodeSubmissionPolicy) for more details.
- While we encourage the release of code and data, we understand that this might not be possible, so [No] is an acceptable answer. Papers cannot be rejected simply for not including code, unless this is central to the contribution (e.g., for a new open-source benchmark).
- The instructions should contain the exact command and environment needed to run to reproduce the results. See the NeurIPS code and data submission guidelines (https://neurips.cc/public/guides/CodeSubmissionPolicy) for more details.
- The authors should provide instructions on data access and preparation, including how to access the raw data, preprocessed data, intermediate data, and generated data, etc.
- The authors should provide scripts to reproduce all experimental results for the new proposed method and baselines. If only a subset of experiments are reproducible, they should state which ones are omitted from the script and why.
- At submission time, to preserve anonymity, the authors should release anonymized versions (if applicable).
- Providing as much information as possible in supplemental material (appended to the paper) is recommended, but including URLs to data and code is permitted.

##### 6. Experimental setting/details

Question: Does the paper specify all the training and test details (e.g., data splits, hyperparameters, how they were chosen, type of optimizer) necessary to understand the results?

Answer: [Yes]

Justification: No model training is performed; the paper evaluates existing models in a zero-shot setting. The folding protocol (MSA generation, no templates, five trunk calls, single diffusion sample, top-ranked structure selection), scoring metric (ipSAE with specified cutoffs), dataset construction filters (sequence identity, length ranges), and simulation setup (1-in-50 trials, 27 targets, 20 trials each) are all described in Sections 2 and 3.

Guidelines:

- The answer [N/A] means that the paper does not include experiments.
- The experimental setting should be presented in the core of the paper to a level of detail that is necessary to appreciate the results and make sense of them.
- The full details can be provided either with the code, in appendix, or as supplemental material.

##### 7. Experiment statistical significance

Question: Does the paper report error bars suitably and correctly defined or other appropriate information about the statistical significance of the experiments?

Answer: [Yes]

Justification: The 1-in-50 screening simulation (Figure 5) reports mean accuracy over 25 independent runs with different random scenario-generation seeds, with *±*2*σ* error bars computed from the sample standard deviation across runs. The pairwise benchmarks operate on the full dataset of 11,247 pairs without random sampling, so error bars do not apply. Appendix J reports bootstrap confidence intervals where subsets of the benchmark are compared.

Guidelines:

- The answer [N/A] means that the paper does not include experiments.
- The authors should answer [Yes] if the results are accompanied by error bars, confidence intervals, or statistical significance tests, at least for the experiments that support the main claims of the paper.
- The factors of variability that the error bars are capturing should be clearly stated (for example, train/test split, initialization, random drawing of some parameter, or overall run with given experimental conditions).
- The method for calculating the error bars should be explained (closed form formula, call to a library function, bootstrap, etc.)
- The assumptions made should be given (e.g., Normally distributed errors).
- It should be clear whether the error bar is the standard deviation or the standard error of the mean.
- It is OK to report 1-sigma error bars, but one should state it. The authors should preferably report a 2-sigma error bar than state that they have a 96% CI, if the hypothesis of Normality of errors is not verified.
- For asymmetric distributions, the authors should be careful not to show in tables or figures symmetric error bars that would yield results that are out of range (e.g., negative error rates).
- If error bars are reported in tables or plots, the authors should explain in the text how they were calculated and reference the corresponding figures or tables in the text.

##### 8. Experiments compute resources

Question: For each experiment, does the paper provide sufficient information on the computer resources (type of compute workers, memory, time of execution) needed to reproduce the experiments?

Answer: [Yes]

Justification: Section 2.3 and appendix L reports that folding all 22,494 complexes across six models required approximately 9,000 GPU-hours on NVIDIA B200 GPUs.

Guidelines:

- The answer [N/A] means that the paper does not include experiments.
- The paper should indicate the type of compute workers CPU or GPU, internal cluster, or cloud provider, including relevant memory and storage.
- The paper should provide the amount of compute required for each of the individual experimental runs as well as estimate the total compute.
- The paper should disclose whether the full research project required more compute than the experiments reported in the paper (e.g., preliminary or failed experiments that didn’t make it into the paper).

##### 9. Code of ethics

Question: Does the research conducted in the paper conform, in every respect, with the NeurIPS Code of Ethics https://neurips.cc/public/EthicsGuidelines?

Answer: [Yes]

Justification: The research uses publicly available patent data and protein databases, evaluates publicly accessible models, and does not involve human subjects, deception, or dual-use concerns beyond standard computational biology research.

Guidelines:

- The answer [N/A] means that the authors have not reviewed the NeurIPS Code of Ethics.
- If the authors answer [No], they should explain the special circumstances that require a deviation from the Code of Ethics.
- The authors should make sure to preserve anonymity (e.g., if there is a special consideration due to laws or regulations in their jurisdiction).

##### 10. Broader impacts

Question: Does the paper discuss both potential positive societal impacts and negative societal impacts of the work performed?

Answer: [N/A]

Justification: The work is a benchmarking study of existing protein structure prediction models on antibody-antigen binding. It does not introduce new models or capabilities that would create direct negative societal impacts. The positive impact of improving antibody discovery pipelines is implicit in the drug discovery context.

Guidelines:

- The answer [N/A] means that there is no societal impact of the work performed.
- If the authors answer [N/A] or [No], they should explain why their work has no societal impact or why the paper does not address societal impact.
- Examples of negative societal impacts include potential malicious or unintended uses (e.g., disinformation, generating fake profiles, surveillance), fairness considerations (e.g., deployment of technologies that could make decisions that unfairly impact specific groups), privacy considerations, and security considerations.
- The conference expects that many papers will be foundational research and not tied to particular applications, let alone deployments. However, if there is a direct path to any negative applications, the authors should point it out. For example, it is legitimate to point out that an improvement in the quality of generative models could be used to generate Deepfakes for disinformation. On the other hand, it is not needed to point out that a generic algorithm for optimizing neural networks could enable people to train models that generate Deepfakes faster.
- The authors should consider possible harms that could arise when the technology is being used as intended and functioning correctly, harms that could arise when the technology is being used as intended but gives incorrect results, and harms following from (intentional or unintentional) misuse of the technology.
- If there are negative societal impacts, the authors could also discuss possible mitigation strategies (e.g., gated release of models, providing defenses in addition to attacks, mechanisms for monitoring misuse, mechanisms to monitor how a system learns from feedback over time, improving the efficiency and accessibility of ML).

##### 11. Safeguards

Question: Does the paper describe safeguards that have been put in place for responsible release of data or models that have a high risk for misuse (e.g., pre-trained language models, image generators, or scraped datasets)?

Answer: [N/A]

Justification: The released dataset contains antibody-antigen sequence pairs extracted from publicly available US patent documents. It does not pose risks for misuse beyond the information already publicly available in the patent literature.

Guidelines:

- The answer [N/A] means that the paper poses no such risks.
- Released models that have a high risk for misuse or dual-use should be released with necessary safeguards to allow for controlled use of the model, for example by requiring that users adhere to usage guidelines or restrictions to access the model or implementing safety filters.
- Datasets that have been scraped from the Internet could pose safety risks. The authors should describe how they avoided releasing unsafe images.
- We recognize that providing effective safeguards is challenging, and many papers do not require this, but we encourage authors to take this into account and make a best faith effort.

##### 12. Licenses for existing assets

Question: Are the creators or original owners of assets (e.g., code, data, models), used in the paper, properly credited and are the license and terms of use explicitly mentioned and properly respected?

Answer: [Yes]

Justification: All models (AlphaFold Multimer, Boltz-2, IntelliFold v2, OpenFold3, Protenix-v1, Protenix-v2), databases (GenBank, SAbDab, OAS, UniProt), and tools (ANARCI, KA-Search, Diamond BLASTP, ColabFold MSAs, ipSAE) are cited with their original publications. The ipSAE tool URL is provided.

Guidelines:

- The answer [N/A] means that the paper does not use existing assets.
- The authors should cite the original paper that produced the code package or dataset.
- The authors should state which version of the asset is used and, if possible, include a URL.
- The name of the license (e.g., CC-BY 4.0) should be included for each asset.
- For scraped data from a particular source (e.g., website), the copyright and terms of service of that source should be provided.
- If assets are released, the license, copyright information, and terms of use in the package should be provided. For popular datasets, paperswithcode.com/datasets has curated licenses for some datasets. Their licensing guide can help determine the license of a dataset.
- For existing datasets that are re-packaged, both the original license and the license of the derived asset (if it has changed) should be provided.
- If this information is not available online, the authors are encouraged to reach out to the asset’s creators.

##### 13. New assets

Question: Are new assets introduced in the paper well documented and is the documentation provided alongside the assets?

Answer: [Yes]

Justification: The paper introduces a new patent-derived antibody-antigen dataset released on Zenodo. The dataset construction pipeline, filtering criteria, and field descriptions are documented in Sections 2.1–2.2 and Appendix A.

Guidelines:

- The answer [N/A] means that the paper does not release new assets.
- Researchers should communicate the details of the dataset/code/model as part of their submissions via structured templates. This includes details about training, license, limitations, etc.
- The paper should discuss whether and how consent was obtained from people whose asset is used.
- At submission time, remember to anonymize your assets (if applicable). You can either create an anonymized URL or include an anonymized zip file.

##### 14. Crowdsourcing and research with human subjects

Question: For crowdsourcing experiments and research with human subjects, does the paper include the full text of instructions given to participants and screenshots, if applicable, as well as details about compensation (if any)?

Answer: [N/A]

Justification: The paper does not involve crowdsourcing or research with human subjects.

Guidelines:

- The answer [N/A] means that the paper does not involve crowdsourcing nor research with human subjects.
- Including this information in the supplemental material is fine, but if the main contribution of the paper involves human subjects, then as much detail as possible should be included in the main paper.
- According to the NeurIPS Code of Ethics, workers involved in data collection, curation, or other labor should be paid at least the minimum wage in the country of the data collector.

##### 15. Institutional review board (IRB) approvals or equivalent for research with human subjects

Question: Does the paper describe potential risks incurred by study participants, whether such risks were disclosed to the subjects, and whether Institutional Review Board (IRB) approvals (or an equivalent approval/review based on the requirements of your country or institution) were obtained?

Answer: [N/A]

Justification: The paper does not involve research with human subjects. Guidelines:

- The answer [N/A] means that the paper does not involve crowdsourcing nor research with human subjects.
- Depending on the country in which research is conducted, IRB approval (or equivalent) may be required for any human subjects research. If you obtained IRB approval, you should clearly state this in the paper.
- We recognize that the procedures for this may vary significantly between institutions and locations, and we expect authors to adhere to the NeurIPS Code of Ethics and the guidelines for their institution.
- For initial submissions, do not include any information that would break anonymity (if applicable), such as the institution conducting the review.

##### 16. Declaration of LLM usage

Question: Does the paper describe the usage of LLMs if it is an important, original, or non-standard component of the core methods in this research? Note that if the LLM is used only for writing, editing, or formatting purposes and does *not* impact the core methodology, scientific rigor, or originality of the research, declaration is not required.

Answer: [Yes]

Justification: Gemini 3 Pro is a core component of the data extraction pipeline, used to parse unstructured patent text and extract antibody-antigen records with up to 36 fields. Its usage is described in Section 2.1, including its role as both extractor and auditor.

Guidelines:

- The answer [N/A] means that the core method development in this research does not involve LLMs as any important, original, or non-standard components.
- Please refer to our LLM policy in the NeurIPS handbook for what should or should not be described.

